# Centrosome Softening As A Mechanical Adaptation For Mitosis

**DOI:** 10.1101/2025.09.09.675178

**Authors:** Júlia Garcia-Baucells, Carlo Bevilacqua, Manuel Rufin, Cornelia Rumpf-Kienzl, Alexandra Zampetaki, Orestis G. Andriotis, Philipp J. Thurner, Robert Prevedel, Sebastian Fürthauer, Alexander Dammermann

## Abstract

Centrosomes are microtubule-organizing centers important for mitotic spindle assembly and chromosome segregation. During mitosis, centrosomes are exposed to mechanical forces via the microtubules they nucleate, yet the material properties underlying their response to these forces remain poorly understood. In this study, we systematically probed the mechanical behavior of *C. elegans* centrosomes, both *in vitro* and *in vivo*. Using microtubule perturbations and quantitative live cell imaging, we found that centrosomes become increasingly deformed during mitosis. Centrosome deformation is independent of cortical pulling forces but instead results from microtubule polymerization within the pericentriolar material. This deformation impacts centrosome size: as microtubule number decreases with cell volume in early cleavage divisions, centrosome size scales proportionately. To directly measure centrosome elasticity, we employed atomic force microscopy (AFM) on isolated centrosomes *in vitro* and Brillouin light scattering microscopy in developing embryos *in vivo*. Both approaches revealed that centrosomes progressively soften during mitosis. Theoretical modeling predicts that softening serves to dampen spindle force fluctuations, helping to protect kinetochore-microtubule interactions and safeguarding chromosome segregation. Further, softening may enhance centrosomal microtubule nucleation capacity, facilitating mitotic spindle assembly, particularly in large early embryonic cells. We propose that centrosome softening is a mechanical adaptation for mitosis that couples microtubule number to centrosome size through force-dependent deformation. This optimally balances two mitotic requirements: the need for robust microtubule nucleation and the ability to withstand spindle forces, thereby ensuring accurate cell division.

## Introduction

Centrosomes are non-membrane-bound organelles which act as the main microtubule-organizing centers in animal cells. As such, they play critical roles in mitosis and male meiosis, facilitating spindle assembly and chromosome segregation and directing both symmetric and asymmetric cell division – processes that are fundamental to organismal development and adult tissue homeostasis (1, 2). Centrosomes are composed of a pair of microtubule-based centrioles surrounded by the pericentriolar material or PCM, a dense accumulation of proteins within which microtubule nucleation and anchoring occurs (3-7).

The PCM undergoes considerable structural reorganization during the cell cycle. During interphase, the PCM forms a thin layer around the centrioles (8, 9). However, as cells enter mitosis, the PCM expands into a matrix-like structure with enhanced microtubule nucleation capacity (9, 10). This expansion is crucial for generating the large number of microtubules required for spindle assembly and cell division. At the end of mitosis, the accumulated PCM is rapidly shed in a manner potentiated by microtubule pulling forces exerted by dynein motors anchored at the cell cortex (11, 12). While centrioles maintain a constant mature size throughout development (13), the mitotic PCM exhibits a pronounced size scaling behavior. This has been particularly well documented for *C. elegans* embryogenesis, where PCM size scales with cell volume (14, 15). PCM size scaling has been proposed to result from limiting amounts of a single centrosomal component (14), with PCM size in turn contributing to regulation of spindle length (16). Remarkably, this scaling appears evolutionarily conserved, with spindle size similarly correlating with cell size across metazoans (17-21).

Mitotic PCM assembly (also referred to as centrosome maturation) has been intensively studied at a molecular level. Central to PCM assembly are scaffolding proteins like SPD-5 in *C. elegans* (22), Cnn in *Drosophila* (23) and CDK5RAP2 in vertebrates (24), which act as a platform for the recruitment of other ‘client’ proteins, including the nucleator γ-tubulin. Pioneering *in vitro* reconstitution experiments have shown that SPD-5 spontaneously self-assembles to form micron-scale scaffolds that can dynamically concentrate client proteins and organize microtubules (25, 26). Similar self-assembly properties have been demonstrated for Cnn and CDK5RAP2, with multivalent interactions between their coiled-coil domains driving multimerization and hence PCM expansion for all three proteins (27-29). Self-assembly *in vitro* is potentiated by the *in vivo* regulators of PCM assembly, most notably PLK-1 (30) and SPD-2/CEP192 (31, 32). PLK-1-mediated phosphorylation enhances scaffold protein oligomerization *in vitro* (25, 27, 28) and is required for PCM recruitment and maintenance *in vivo* (33, 34). In addition to promoting scaffold assembly, PLK-1 also regulates client protein recruitment (35). SPD-2/CEP192 also promotes PCM expansion, both directly through promoting scaffold polymer self-assembly (25) and indirectly through recruiting PLK-1 to the centrosome (31, 34, 36, 37).

However, centrosomes are not merely molecular hubs – they also function as mechanical elements that must withstand substantial forces acting on the microtubules they nucleate. This mechanical role is readily apparent from the rapid movement of centrosome remnants towards the cortex following laser ablation-induced fragmentation in late mitosis (38). At earlier stages of mitosis, centriole ablations similarly trigger rapid, microtubule-dependent PCM disassembly (34). The latter finding not only serves to illustrate the magnitude of the forces acting on the PCM, but also the stabilizing effect of the centrioles, which is particularly remarkable given their small size compared to the order of magnitude larger mitotic PCM they anchor, underscoring the importance of their mechanical coupling.

The ability to withstand forces, therefore, is fundamental to centrosome function in cell division as well as in other cellular processes, such as cell migration (39). Yet, in contrast to the molecular mechanisms underlying PCM assembly, the mechanical behavior of the PCM remains poorly understood. In fact, the material state of the PCM itself remains a subject of active debate (40). One model originally derived from *in vitro* reconstitution studies describes the PCM scaffold as a phase-separated biomolecular condensate (26, 41). According to this view, an initial liquid-like state accounts for its isotropic growth (42), while subsequent hardening into a more solid phase could explain its capacity to withstand mitotic forces. The alternative view, derived from ultrastructural analysis of intact centrosomes depicts the PCM as a loose meshwork of fibers permeated by cytoplasm which remains solid throughout the cell-cycle (43, 44). What is clear is that centrosome mechanical properties are not constant throughout the cell cycle. Recent work using optical induction of cytoplasmic flows to mechanically probe centrosomes has shown that the PCM at anaphase/telophase undergoes a transition from a deformation-resistant state to a weaker material that is susceptible to deformation and rupture (45). However, these methods could not probe earlier mitotic stages, when centrosomes are most critical for spindle assembly and stability (46).

To address centrosome material properties more comprehensively, we here used a combination of quantitative live cell imaging, atomic force microscopy (AFM) of isolated centrosomes and *in vivo* Brillouin microscopy to examine centrosomes throughout mitosis using the *C. elegans* early embryo as an experimental model. Our results demonstrate that centrosomes progressively soften during mitosis, with their size being determined primarily by internal microtubule polymerization forces. We propose that this mechanical adaptation is a critical regulator of microtubule nucleation capacity and spindle stability, ensuring accurate chromosome segregation.

## Results

### The PCM Scaffold Progressively Deforms During Mitosis

The PCM is a dynamic structure critical for centrosome function in cell division, but how it responds to mechanical forces remains unclear. We therefore asked whether the PCM is deformed by microtubule-dependent forces and how its material properties influence this response. To address this, we used nocodazole to depolymerize microtubules in *C. elegans* embryos and analyzed the resulting changes in PCM size using endogenously GFP-tagged SPD-5 as a marker of the PCM scaffold (Fig. 1A). To ensure consistent drug entry, we permeabilized embryos using *perm-1* RNAi (47), while simultaneously inhibiting the spindle checkpoint with *mad-2*^*mdf-2*^ RNAi to prevent mitotic arrest (48). Embryos were imaged by spinning disk confocal microscopy before and after acute nocodazole treatment at nuclear envelope breakdown (NEBD).

**Figure 1.**
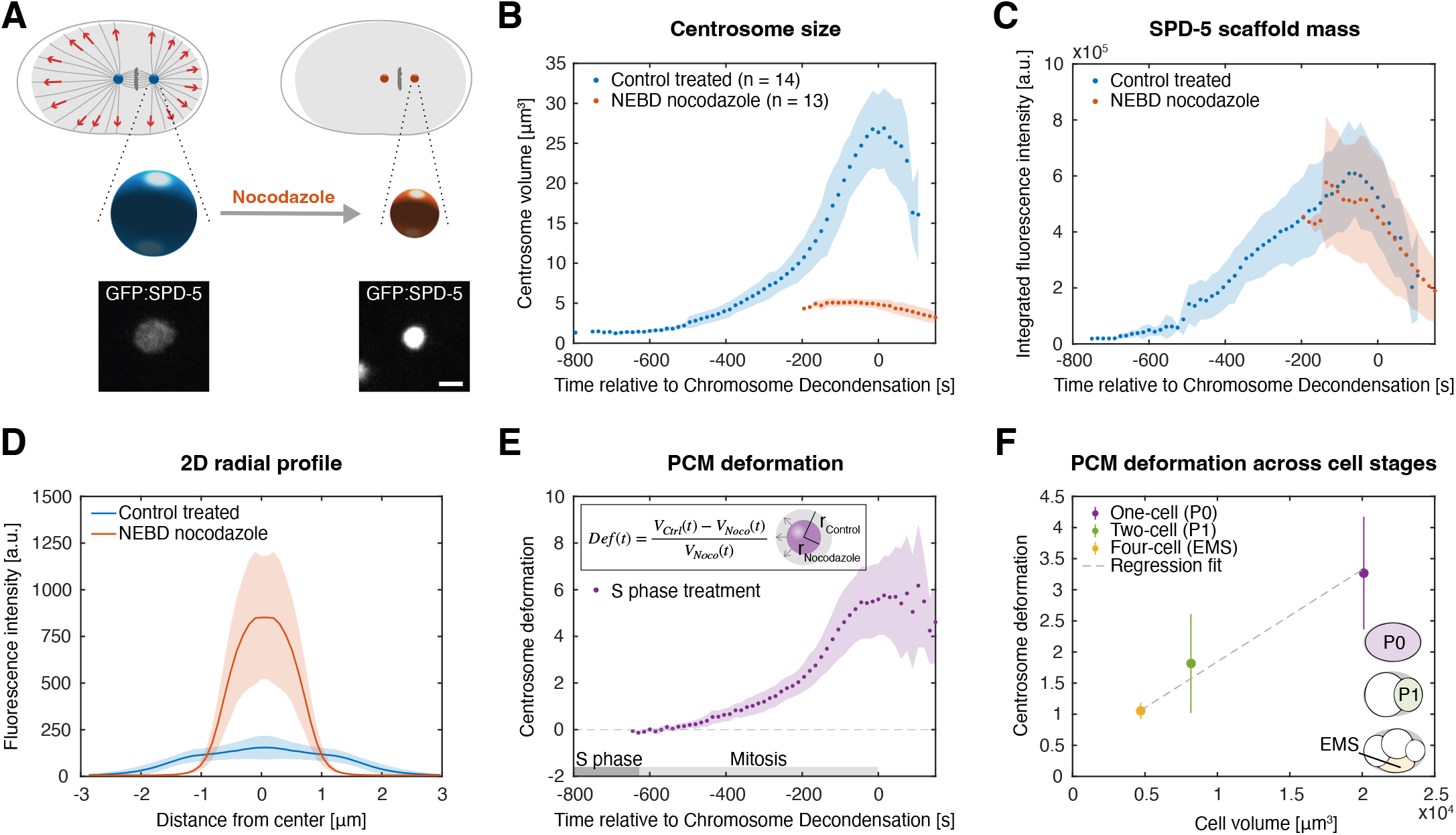
Microtubule-dependent PCM deformation increases during mitosis and scales with cell volume in early *C. elegans* embryos. (A) Permeabilized and spindle assembly checkpoint-inhibited *perm-1;mad-2(RNAi)* embryos were treated with nocodazole in S phase or at nuclear envelope breakdown (mitosis) and the effect on PCM scaffold size was monitored using GFP:SPD-5. The mitotic PCM visibly contracts upon nocodazole treatment. Centrosomes at cytokinesis onset are shown as examples of control and nocodazole-treated conditions. Spheres represent centrosome volume to scale for comparison. Scale bar: 1 µm. (B) Centrosomes expand dramatically in size upon entry into mitosis, but this expansion is largely abolished by nocodazole treatment. See Table S2 for a timeline of events in the first embryonic cell cycle. (C) Nocodazole treatment does not result in a loss of PCM scaffold. (D) 2D radial profiling of GFP:SPD-5 fluorescence intensities at the centrosome midplane in nocodazole-treated mitotic embryos shows that the PCM polymer increases in density close to the centrioles compared to controls. (E) Microtubule-dependent PCM deformation increases progressively during mitosis but is negligible in S phase centrosomes. Nocodazole treatment in S phase was used as a baseline (control n = 14, nocodazole n = 9), data are from Fig. S1D. (F) Mitotic PCM deformation (measured from -100 to -50 s relative to chromosome decondensation; dashed lines in Fig S1J, using nocodazole treatment at NEBD as the baseline, data are from Fig. S1J) scales with cell volume across one-, two- and four-cell stage embryos. Points in B-F indicate the mean of anterior and posterior centrosomes (averaged per embryo) across embryos; error bars or shading represent the associated standard deviation.

To reliably quantify PCM size, we developed an image analysis pipeline combining two independent methods for centrosome size measurement in the 2D midplane: 1) intensity-based segmentation using Otsu thresholding (49), and 2) radial intensity profiling (50) (Fig. S1A). For the radial intensity profiling method, concentric ellipses were drawn from the centrosome center and the radius was derived from the full-width half-maximum (FWHM) of the Gaussian-fitted intensity profile. The close agreement between the two methods (Fig. S1B) validated our approach. We extended this analysis to 3D by applying Otsu thresholding to the full centrosome volume to determine the equidistant 3D radius. This identified a systematic 0.3 µm offset between these 3D radii and the 2D midplane measurements (Fig. S1C). As this offset was size-independent, we used intercept-corrected 2D data for all subsequent analyses, calculating PCM volume from the 2D radii under the assumption of spherical geometry (see Methods).

To understand how PCM material properties might affect deformation, we envisioned three potential scenarios: 1) if the PCM were a rigid non-deformable solid, neither its volume nor its shape would be affected by microtubule depolymerization; 2) if it were an incompressible viscous liquid, its volume would be maintained although its shape might change; and 3) if it were an elastically deformable solid, its shape and volume would be altered without material loss upon force removal. Using our analysis pipeline, we found that nocodazole-treated centrosomes retained the same amount of GFP:SPD-5 PCM scaffold polymer as controls (p = 0.072, n.s.), yet were significantly smaller in size (volume reduction of 80.3%, p = 4.9e-90) (Fig. 1B, C). Consistent with this, radial fluorescence intensity profiles revealed a higher scaffold density in nocodazole-treated centrosomes (Fig. 1D). These results rule out scenarios 1) and 2) (both of which require constant volume), but are in line with scenario 3). Given that the PCM’s relaxation time is approximately 100 s, as suggested by Paulin et al. (51), which is comparable to our experimental timescale, this justifies treating the PCM scaffold as an elastically deformable solid for the purposes of the analysis in this manuscript. Additionally, the accumulation of a similar polymer mass in both control and nocodazole conditions indicates that PCM assembly occurs largely independently of microtubule-mediated transport, consistent with previous findings (10). Together, these results demonstrate that microtubule-generated forces stretch the PCM scaffold during mitosis, and that on short timescales, the PCM behaves as an elastically deformable solid, expanding to a multiple of its relaxed size.

Building on the idea that microtubule forces stretch the mitotic PCM scaffold, we next asked whether this deformation remains constant or changes dynamically across the cell cycle. To test this, we treated *perm-1;mad-2* RNAi one-cell stage embryos with nocodazole at S phase and quantified centrosome strain by calculating the degree of PCM deformation per unit PCM (*Volume*_*Control*_ − *Volume*_*Nocodazole*_)/*Volume*_*Nocodazole*_ over time. Our analysis revealed that PCM deformation is not static but develops progressively: while S phase centrosomes are not measurably deformed, deformation first becomes detectable at mitotic onset and increases throughout mitosis (Fig. 1E, S1D). The complete absence of deformation in S phase centrosomes – despite active microtubule nucleation and centrosome movements during this stage – demonstrates that microtubule forces only deform the centrosome after the onset of maturation. This mitotic deformation has functional consequences for PCM growth. As predicted by the strain-limited polymer growth model (51), we found that S phase-treated centrosomes, which are not stretched, ultimately reached smaller final sizes than those treated at NEBD, which were allowed to grow under tension (Fig. S1K). This size difference arises because, on the longer time scales that govern centrosome growth, centrosome strain limits the incorporation of new scaffold material (51).

Given the proportional scaling between cell size and mitotic centrosome volume during *C. elegans* embryogenesis reported by Decker et al. (14), we asked whether centrosome deformation changes across successive cell divisions. To address this, we extended our nocodazole treatment experiments to two- and four-cell stage embryos and quantified centrosome deformation as described above (Fig. S1E-I). In control embryos, centrosome volume scaled with cell size, recapitulating the findings of Decker et al. (Fig. S1K). The limiting component model proposes that centrosome size is set by a limited pool of centrosomal components, the amount of which is determined by cell volume. According to this model, with decreasing cell volume and consequently less PCM material, we would expect constant or even increased deformation depending on the extent to which cortical force generators are also reduced. However, what we actually observed was the opposite: deformation decreased with each division as cell volume halved (Fig. 1F, S1J,K). This result is inconsistent with the limiting component model. Indeed, nocodazole-treated centrosomes reached approximately the same ultimate size at one-, two- and four-cell stages (Fig. S1K,L), demonstrating that mitotic size is not dictated by a specific centrosomal structural component.

In summary, these results indicate that mitotic centrosomes behave as elastically deformable solids, undergoing progressive deformation during mitosis. This deformation scales with cell size, with centrosomes in two- and four-cell stage embryos exhibiting proportionally less PCM stretch than those at the larger one-cell stage. These findings challenge a model in which centrosome size is determined by a limiting centrosomal component and instead point to cell cycle stage-specific, mechanical regulation of PCM deformation.

### Cortical Forces Do Not Significantly Contribute To Centrosome Deformation

Having established that centrosome deformation scales with cell size and becomes more pronounced during mitosis, we next asked whether dynein-mediated cortical pulling forces are responsible for this deformation. To test this, we modulated cortical forces without altering microtubule numbers or dynamics, increasing forces via *efa-6* RNAi (which extends microtubule residency time at the cell cortex (52)) or decreasing forces via *gpr-1/2* RNAi (disrupting the cortical dynein complex (38, 53, 54)) (Fig. 2A). We first confirmed successful perturbation of cortical pulling forces for each condition. In *efa-6* RNAi embryos, increased cortical pulling forces were evidenced by 1) significantly longer final spindle lengths (p = 0.007; embryos with transversely aligned spindles excluded from ANOVA analysis; Fig. S2A,B), 2) transverse spindle alignment corrected by spindle rotation during cytokinesis in 5/14 embryos, reminiscent of the strong microtubule-dependent cortical pulling forces counteracting long-axis finding observed in the 2-cell stage AB cell (55), and 3) centrosome pronuclear detachment and increased centrosome separation during pronuclear migration (Fig. S2A), consistent with prior reports (52). Conversely, *gpr-1/2* RNAi reduced cortical forces, as evidenced by significantly shorter spindles (p = 1.9e-08; Fig. S2A,B) and a loss of anaphase spindle oscillations (Fig. S2C).

**Figure 2.**
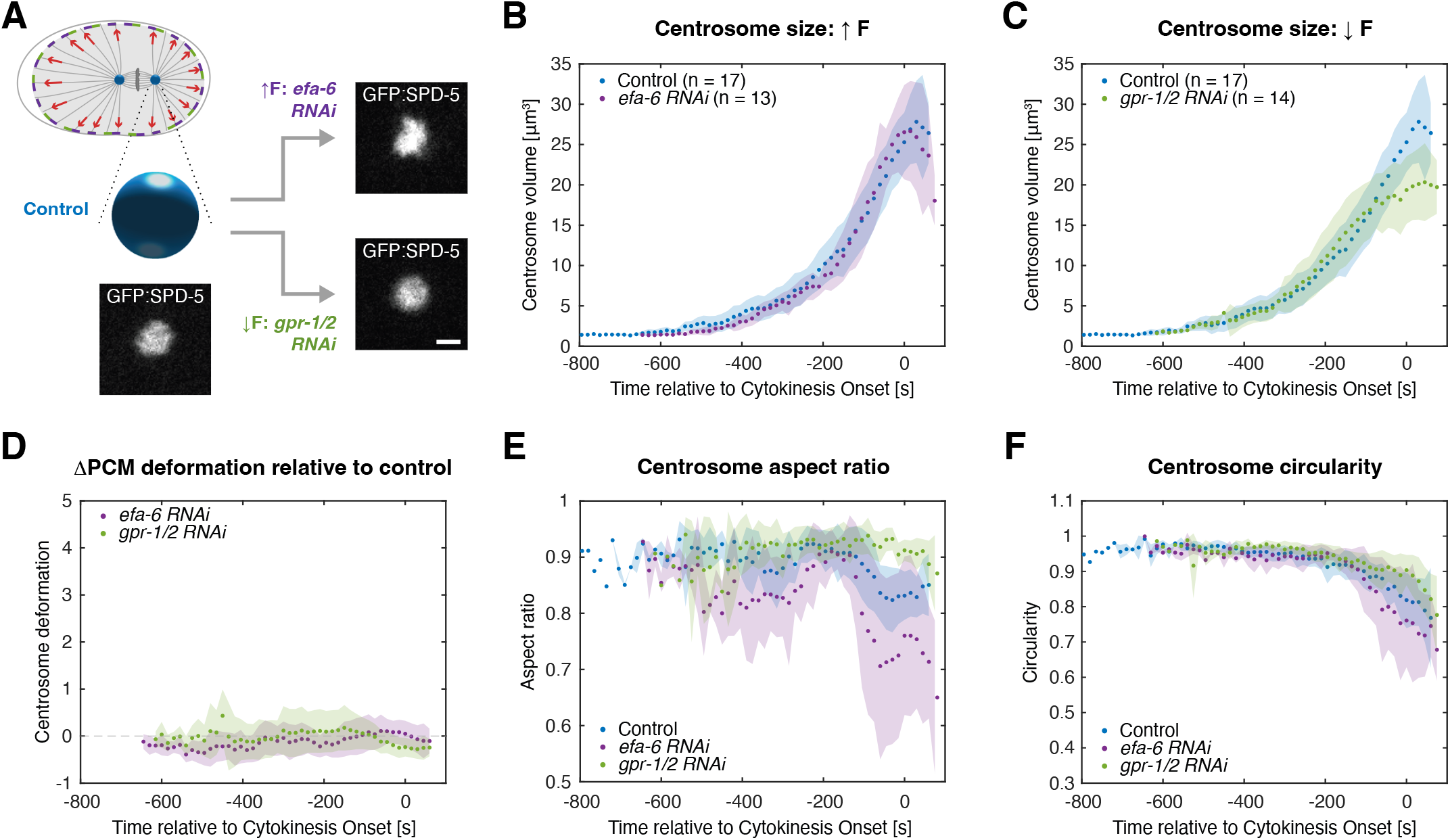
Cortical force modulation fails to alter centrosome deformation. (A) Increasing cortical pulling forces by *efa-6(RNAi)* or inhibiting forces by *gpr-1/2(RNAi)* does not visibly affect mitotic centrosome size, as assessed using GFP:SPD-5. Examples of centrosomes at anaphase onset are shown across conditions for comparison. Scale bar: 1 µm. (B, C) Quantitation of centrosome size during the first embryonic cell cycle upon EFA-6 (B) or GPR-1/2 (C) depletion reveals no significant differences to controls during centrosome maturation, although decreased forces impair PCM disassembly at mitotic exit, as previously reported (11). (D) ΔPCM deformation relative to controls, quantified as (*Volume*_*Control*_ − *Volume*_*RNAi*_)/*Volume*_*RNAi*_, is unchanged in EFA-6 or GPR-1/2-depleted embryos, data are from Figs. 2B and 2C. (E, F) Cortical forces do distort centrosomes from a perfectly spherical shape, with EFA-6-depleted centrosomes being less round and GPR-1/2-depleted centrosomes more so, as assessed using aspect ratio (E) and circularity (F). In B-F, points indicate the mean of anterior and posterior centrosomes (averaged per embryo) across embryos; shading represents the associated standard deviation.

Despite these clear effects on cortical forces, neither perturbation significantly impacted centrosome volume. *efa-6* RNAi led to no detectable change in centrosome volume compared to controls (p = 0.191; Fig. 2B), while *gpr-1/2* RNAi resulted in only a slight reduction (p = 0.023; Fig. 2C) that occurred post-anaphase onset, consistent with known delays in PCM disassembly in this condition (11, 12, 14). Indeed, our normalized deformation analysis (*Volume*_*Control*_ − *Volume*_*RNAi*_)/*Volume*_*RNAi*_ revealed Δ deformation values near zero for both perturbations throughout mitosis (Fig. 2D), demonstrating no measurable effect on centrosome expansion. However, cortical force alterations did induce changes in centrosome morphology. Aspect ratios deviated from 1 (away from a perfect circle) during cell cycle phases characterized by strong imbalanced cortical forces (56) (Fig. 2E) and circularity progressively decreased during mitosis, with *efa-6* RNAi embryos displaying more distortion than controls and *gpr-1/2* RNAi showing less (Fig. 2F). These results show that while cortical forces apply anisotropic stresses that distort PCM shape, centrosomes maintain overall volume.

To understand this force-shape relationship, we investigated how microtubule forces spatially pattern centrosome deformation given the observed shape anisotropy. We first asked whether centrosome deformation is spatially biased (e.g., aligned with the spindle axis) or occurs randomly. To test this, we quantified the orientation of centrosome deformation in control embryos by measuring the angle between the long axis of fitted centrosome ellipses and the spindle axis (defined by anterior/posterior centrosome positions, Fig. S2D). Interestingly, deformation primarily aligned along the spindle axis throughout mitosis (angles close to 0°, Fig. S2E,F), although the posterior centrosome to a lesser extent also showed distortion perpendicular to the spindle axis (angles near ±90°, Fig. S2F), likely induced by the larger number of force generators at the posterior cortex (38). This suggests microtubule-dependent cortical pulling forces propagate along the spindle to anisotropically deform centrosomes.

To confirm that anisotropy emerges from spindle-associated microtubules rather than local polymerization effects, we next asked if deformation direction correlates preferentially with microtubules extending toward the cortex or the central spindle or short microtubules enriched at the centrosome boundary. We imaged embryos expressing GFP-tagged β-tubulin and quantified tubulin intensity in two regions: 1) the centrosome boundary, which contains 50% polymerized tubulin (7), and 2) a distal ring 3-5 µm from the centrosome center (Fig. S2G). While centrosome boundary tubulin fluorescence levels were homogeneous for both anterior and posterior centrosomes (Fig. S2H,J), distal regions showed a pronounced enrichment toward the central spindle (Fig. S2I,K), precisely aligning with the centrosome deformation axis.

Together, these results demonstrate that centrosome deformation is primarily spindle-aligned, with microtubule organization patterns suggesting spindle-associated microtubules as the source of centrosome shape deformation forces. However, critically, because neither cortical force enhancement (*efa-6* RNAi) nor reduction (*gpr-1/2* RNAi) altered the magnitude of mitotic centrosome deformation, only its directionality, we conclude that cortical forces are not the primary driver of PCM volumetric deformation.

### Mitotic Centrosomes Are Primarily Deformed By Microtubule-Generated Forces Within The PCM Scaffold, Which Define Centrosome Size During Mitosis

Having ruled out cortical forces as the principal driver of mitotic centrosome volumetric deformation, we sought to identify the origin of the pronounced size difference between control and nocodazole-treated centrosomes. We hypothesized that centrosome size may be controlled by the number of microtubules embedded within the PCM scaffold. To test this, we increased the number of astral microtubules using RNAi against *klp-7*, which encodes the *C. elegans* ortholog of the kinesin-13 microtubule depolymerase MCAK (57) (Fig. 3A). We validated successful knockdown in GFP:histone H2B; GFP:β-tubulin embryos, which exhibited both increased centrosomal microtubule levels compared to controls and the expected spindle breakage during anaphase (57-59) (Fig. S3A). *klp-7* RNAi further led to elevated inter-centrosome distances in embryos expressing GFP:SPD-5 at the one-cell stage (Fig. S3B), while reducing inter-centrosome distances prior to NEBD in two-cell stage P1 cells (Fig. S3C), as reported previously (60).

**Figure 3.**
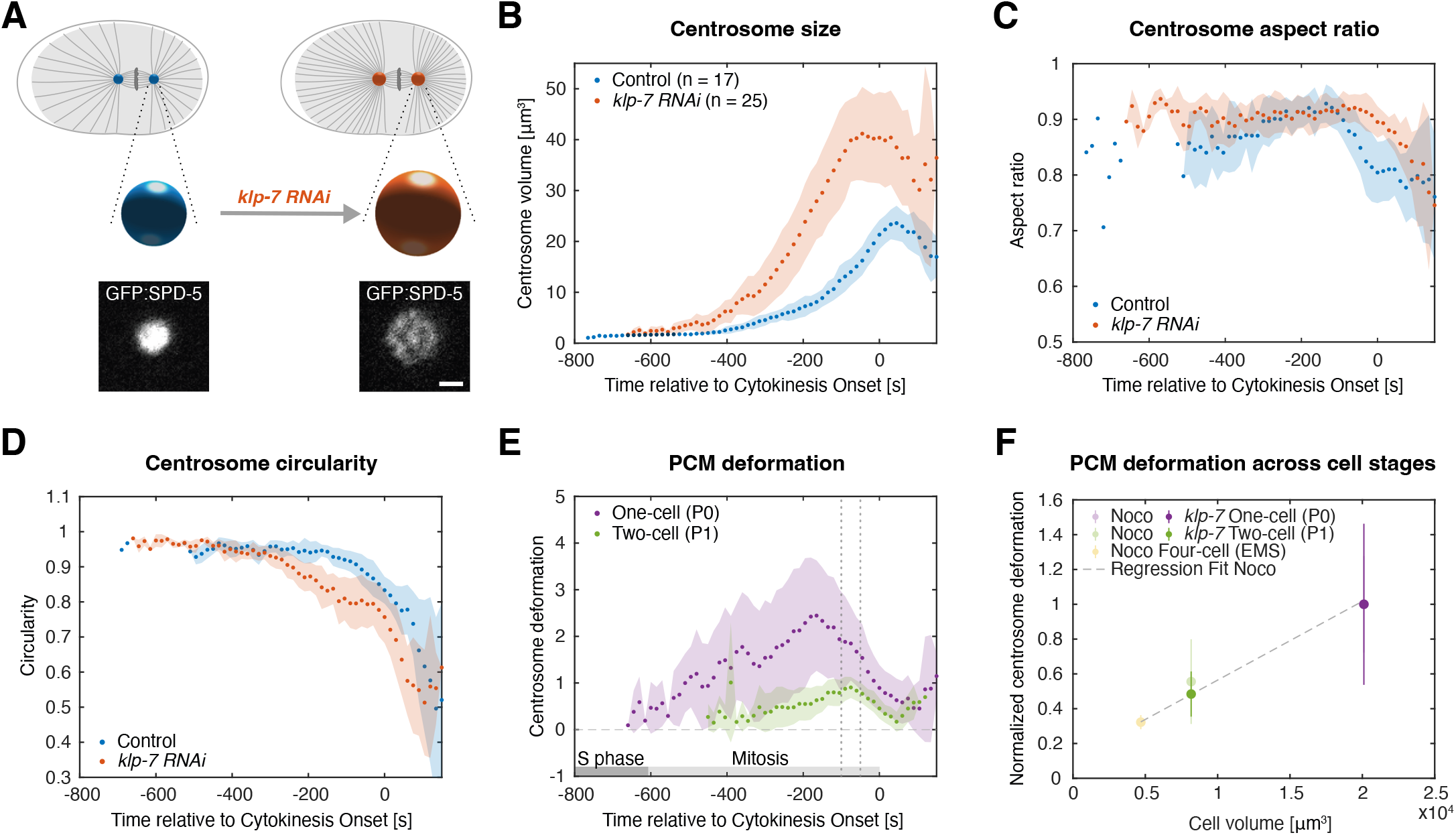
Microtubule-generated forces control centrosome size in mitosis. (A) Increasing centrosomal microtubule numbers via *klp-7(RNAi)* substantially increases centrosome size, as assessed using GFP:SPD-5. Control and *klp-7* RNAi treated centrosomes at metaphase onset are shown, with spheres representing centrosome volumes to scale. Scale bar: 1 µm. (B) Quantification of centrosome size during the first embryonic cell cycle upon *klp-7(RNAi)* confirms increased dimensions throughout mitosis. (C, D) KLP-7 depletion uniformly distorts centrosomes, as assessed using aspect ratio (largely unchanged; C) and circularity (decreased; D). (E) ΔPCM deformation relative to controls, calculated as, (*Volume*_*klp* −7 *RNAi*_ − *Volume*_*Control*_)/*Volume*_*Control*_, reveals stronger effect in one-cell stage (P0) compared to two-cell stage (P1) *klp-7* (RNAi) embryos (2-fold reduction in P1). Data from Fig. 3B (one-cell) and S3D (two-cell). (F) In *klp-7* RNAi embryos, mitotic PCM deformation (measured during -100 to -50 s relative to cytokinesis onset; see dotted lines in E) shows reduced deformation that scales with decreasing cell volume in one- and two-cell stage embryos. Deformation for both *klp-7(RNAi)* (from E) and nocodazole treatment (from Fig. 1F) was normalized to the maximum value in one-cell embryos. The normalized deformation from both perturbations overlaps, revealing a shared linear scaling relationship. In B-F, points represent the mean of anterior/posterior centrosomes (averaged per embryo) across embryos, with error bars or shading indicating standard deviation.

Consistent with our hypothesis, *klp-7* RNAi in one-cell embryos expressing GFP:SPD-5 significantly enlarged mitotic centrosomes compared to controls (volume increase: 105.4%, p = 7.2e-33; Fig. 3B), demonstrating that microtubules within the PCM substantially influence centrosome size during mitosis. The magnitude of this effect is much larger than expected from passive microtubule volume occupancy alone. In control centrosomes, microtubules account for only ∼3% of the centrosome volume at metaphase ((7); see Methods for calculation). If microtubule volume occupancy were the primary determinant of centrosome size, we would expect centrosomes to be at most ∼3% smaller upon microtubule depolymerization, with *klp-7 RNAi* resulting in a similarly minor increase (57, 61). The magnitude of the observed effects instead suggest that microtubules exert active forces, potentially via pushing on the scaffold polymer as they are nucleated and polymerize within the PCM.

Building on our finding that centrosome size increases with microtubule number, we hypothesized that microtubule-induced deformation would increase during mitosis, as this is when centrosomal microtubule nucleation peaks (57, 62). We tested this by quantifying relative deformation using the metric, (*Volume*_*klp* −7 *RNAi*_ − *Volume*_*Control*_)/*Volume*_*Control*_ over time. This analysis revealed that centrosomes progressively deformed during mitosis, with Δ deformation peaking during spindle assembly while S-phase centrosomes remained mostly undeformed (Fig. 3E). These results demonstrate that microtubule-dependent centrosome deformation is mitosis-specific.

We next asked whether *klp-7 RNAi*, like *efa-6* and *gpr-1/2 RNAi*, alters centrosome morphology. We quantified centrosome shape by measuring both aspect ratio and circularity. Surprisingly, while *klp-7 RNAi*-treated centrosomes maintained significantly rounder aspect ratios than controls despite their increased microtubule density (Fig. 3C), their circularity decreased during mitosis (Fig. 3D). These results suggest that aspect ratio primarily reflects external cortical forces, while circularity captures the combined effects of internal microtubule polymerization forces and external cortical pulling forces.

Given that centrosomes in later-stage embryos show reduced deformation in nocodazole (Fig. 1F), we asked whether two-cell stage centrosomes – which likely contain half the microtubule number due to volume-scaling (63) – follow the same deformation-volume scaling relationship. If deformation depends linearly on microtubule number, *klp-7* RNAi-treated two-cell stage centrosomes should exhibit proportionately reduced deformation compared to their one-cell stage counterparts. We tested this by quantifying deformation in *klp-7* RNAi-treated two-cell stage embryos (Fig. S3D). Indeed, two-cell stage centrosomes displayed 2-fold less deformation than equivalently treated one-cell stage centrosomes (Fig. 3E), matching their cell volume reduction. Strikingly, when normalized, the *klp-7(RNAi)*-induced deformation perfectly matched the trend of the nocodazole-derived deformation, with the data from both perturbations overlapping (Fig. 3F). This linear scaling demonstrates that centrosome deformation – and consequently mitotic centrosome size – depends directly on the microtubule number set by cell volume (Fig. 3F).

In summary, these results demonstrate that centrosomes are predominantly deformed by PCM-embedded microtubules and that this microtubule-dependent deformation is mitosis-specific, scaling linearly with microtubule number and cell volume. Collectively, these results support a mechanical regulation model in which centrosome size is controlled by microtubule polymerization forces within the PCM scaffold.

### Centrosome Stiffness By AFM Varies With Centrosome Size, With Larger Mitotic Centrosomes Being Softer

The observed linear scaling between centrosome deformation and microtubule number led us to consider two potential explanations for their progressive deformation during mitosis. First, the PCM might maintain constant mechanical properties throughout the cell cycle, with deformation being determined solely by the number of microtubules embedded within the PCM scaffold. Alternatively, the centrosome could undergo softening during mitosis, potentially facilitating increased microtubule incorporation and consequent deformation. To distinguish between these possibilities and directly assess centrosome mechanical properties, we applied AFM to centrosomes isolated from *C. elegans* early embryos expressing GFP:SPD-5 using a protocol adapted from *Drosophila* (64, 65) (see Methods, Fig. 4A, B). Centrosomes were shown to be functional by their capacity to nucleate and organize microtubules (Fig. 4C).

**Figure 4.**
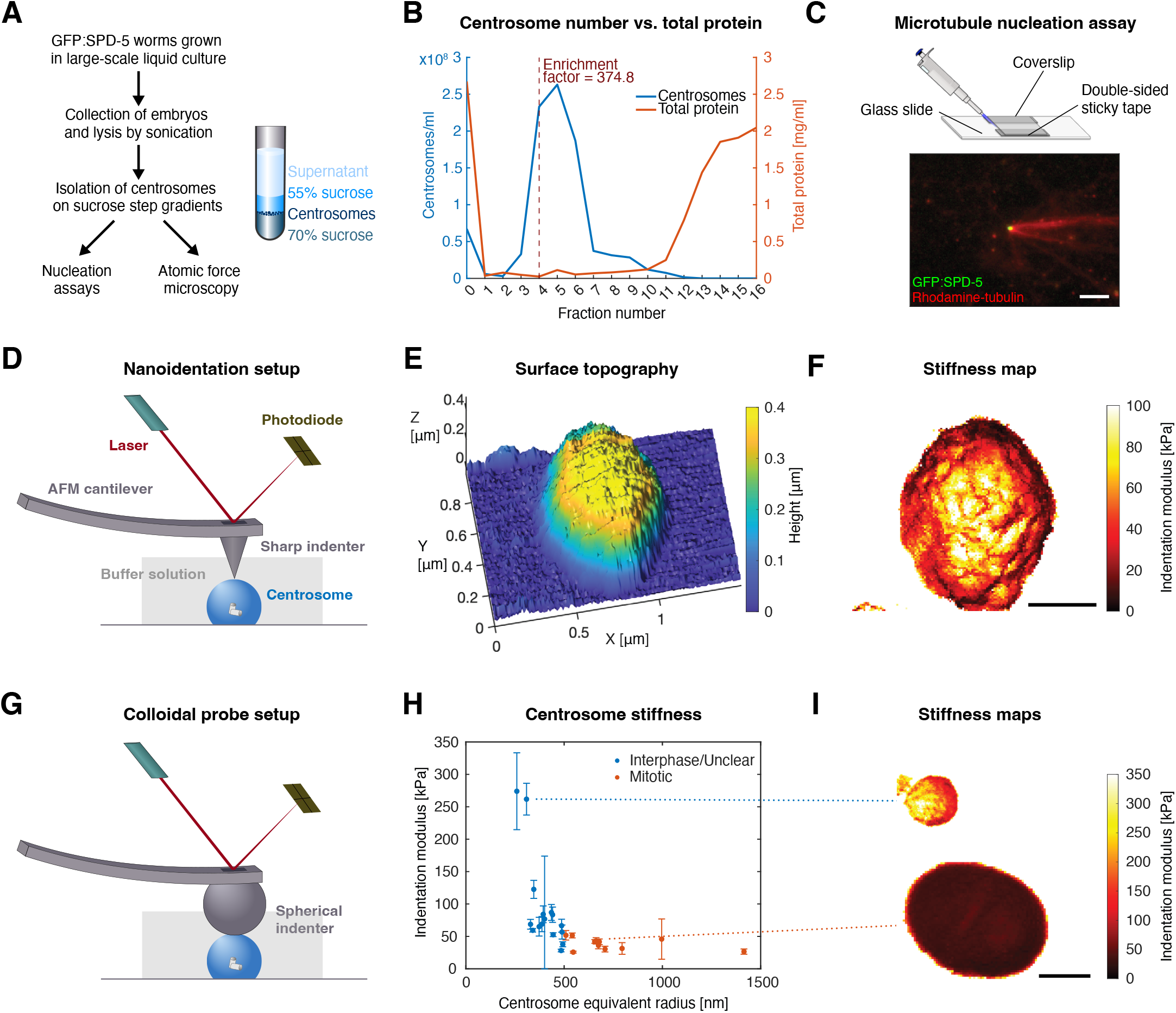
AFM of isolated centrosomes reveals size-dependent changes in stiffness. (A) Protocol for isolating centrosomes from *C. elegans* embryos using sucrose step gradients. (B) Centrosome enrichment, calculated as the number of GFP positive centrosomes per ml in a representative preparation. The most concentrated fractions (here, fractions 4-6) were used for subsequent experiments. (C) Isolated centrosomes retain microtubule nucleation potential. Scale bar: 5 µm. (D) Setup for high-resolution AFM scanning using a nanoindentation probe. (E, F) Surface topography (E) and micromechanical stiffness (F) of isolated centrosomes probed by nanoindentation. Scale bar: 0.5 µm. (G) Setup for high-throughput AFM scanning using colloidal bead probe. (H) Colloidal probe measurements reveal that centrosome stiffness decreases with size, with mitotic centrosomes >0.5 µm being softer than their smaller counterparts (n = 27 centrosomes). (I) Example of stiffness measurements for large and small centrosomes. Scale bar: 1 µm.

Initial mechanical characterization was performed using high-resolution AFM scanning in physiological buffer (BRB80). Centrosomes were first localized via fluorescence imaging before AFM interrogation. We employed the instrument’s quantitative imaging (QI) mode with a sharp cantilever (nominal tip radius: 2 nm) (Fig. 4D, Fig. S4A), which provides both height topography mapping and spatially resolved nanoscale indentation modulus measurements. From the topographical maps, we observed that centrosomes visibly flatten against the glass surface (Fig. 4E). High-resolution scans further revealed that the PCM surface displays considerable nanoscale mechanical heterogeneity, with areas of high stiffness separated by low stiffness grooves whose biological significance remains unclear (Fig. 4F).

While this high-resolution approach provided excellent spatial detail, its low throughput limited statistical analysis of centrosome stiffness variations. To enable high-throughput mechanical measurements of multiple centrosomes, we therefore implemented colloidal probe force mapping (66) (Fig. 4G, Fig. S4A). This approach offers greater stability for studying soft biological samples albeit at the expense of spatial resolution. For probe fabrication, we attached a 2 μm diameter colloidal bead to a tipless cantilever. To ensure measurement accuracy, we restricted indentation modulus analysis to flat PCM regions to minimize topographical artifacts (Fig. S4B) and applied curvature correction for spherical geometry (see Methods). Only high-quality Hertz fits meeting strict criteria (Predictive R^2^ > 0.96; R^2^ > 0.99) were included in the final mechanical analysis (Fig. S4C-E). The centrosomes in our sample derive from blastomeres at different stages of early embryonic development and different cell cycle stages. In the absence of molecular markers to distinguish between interphase and mitotic centrosomes, we used size as a proxy, classifying centrosomes >0.5 μm in radius as mitotic and <0.5 μm as interphase or early mitotic based on measurements from nocodazole-treated embryos at NEBD across cell stages (Fig. S1L). Our measurements revealed a striking size-dependent mechanical transition (n = 27; distance correlation: 0.6451; p-value: 0.0025). Centrosomes greater than 0.5 μm in radius and hence mitotic in origin exhibited a 2.4-fold lower stiffness compared to their smaller counterparts (p = 3.5e-04; Fig. 4H,I, S4F). The smaller centrosome population displayed a gradient of stiffness values, likely reflecting their heterogeneous composition of interphase centrosomes and early mitotic centrosomes from later embryonic divisions. While we obtained differential compression across samples (with some centrosomes exceeding 40% compression), a greater degree of compression did not correlate with stiffness extremes (Fig. S4G), ruling out compression artifacts as the primary driver of the size-stiffness relationship. To address potential substrate effects – where smaller centrosomes might appear artificially stiff due to their finite thickness – we combined thin film and curvature corrections. These modifications reduced but did not eliminate the nonlinear size-stiffness relationship (distance correlation: 0.4929; p-value: 0.0455, Fig. S4H), confirming that larger mitotic centrosomes are indeed softer.

It is worth noting that, since softening persisted in microtubule-free isolated centrosomes, it represents an intrinsic structural adaptation of the PCM scaffold to mitosis rather than a microtubule-dependent effect. Taken together, these findings provide strong evidence for direct, cell cycle-regulated modulation of PCM material properties.

### Brillouin Microscopy Reveals Progressive Mitotic Softening Of Centrosomes In Live *C. elegans* Embryos

While our AFM measurements of isolated centrosomes revealed size-dependent stiffness variations, with larger, mitotically derived centrosomes being softer, the developmental heterogeneity of these centrosomes prevented precise cell-cycle staging. To address this limitation and assess centrosome mechanical properties directly inside the living embryo, we turned to Brillouin light scattering microscopy, a contact-free optical method to assess material properties (67-69). Brillouin microscopy measures the Brillouin frequency shift, which occurs when light interacts with spontaneous acoustic waves (phonons) emanating from within the sample. The shift these phonons impart is proportional to their velocity and hence provides information about the viscoelastic properties of the material. From the Brillouin shift, the longitudinal modulus – a direct measure of material stiffness – can be calculated if the refractive index and mass density are known (67-69).

Examining mitotic one-cell stage embryos using fluorescence-correlated Brillouin microscopy in a strain expressing GFP:SPD-5 and GFP:histone H2B to label centrosomes and chromosomes, respectively, we found that the spindle space exhibited considerably lower Brillouin shift values than the surrounding cytoplasm (Fig. 5A). Given that embryos contain abundant yolk granules that are rich in lipids (resulting in high refractive index but low mass density), and because the Brillouin shift is proportional to the ratio of refractive index and mass density, we initially questioned whether these high cytoplasmic Brillouin shifts were artifactual. To test this, we used *rme-2* mutants, which are deficient in yolk endocytosis (70) (Fig. S5A). We found no differences in either refractive index, as measured by holotomography (Fig. S5B), or Brillouin shift (Fig. S5C) compared to controls. The high Brillouin shift values in the cytoplasm are therefore likely not artifacts caused by lipid content, but reflect macromolecular crowding in the cytoplasm. In contrast, the spindle space is protected by endomembranes derived from the endoplasmic reticulum and nuclear envelope, which act as a size-selective barrier that excludes large macromolecules (71-75). This exclusion likely contributes to the spindle’s lower density and apparent ‘softness’.

**Figure 5.**
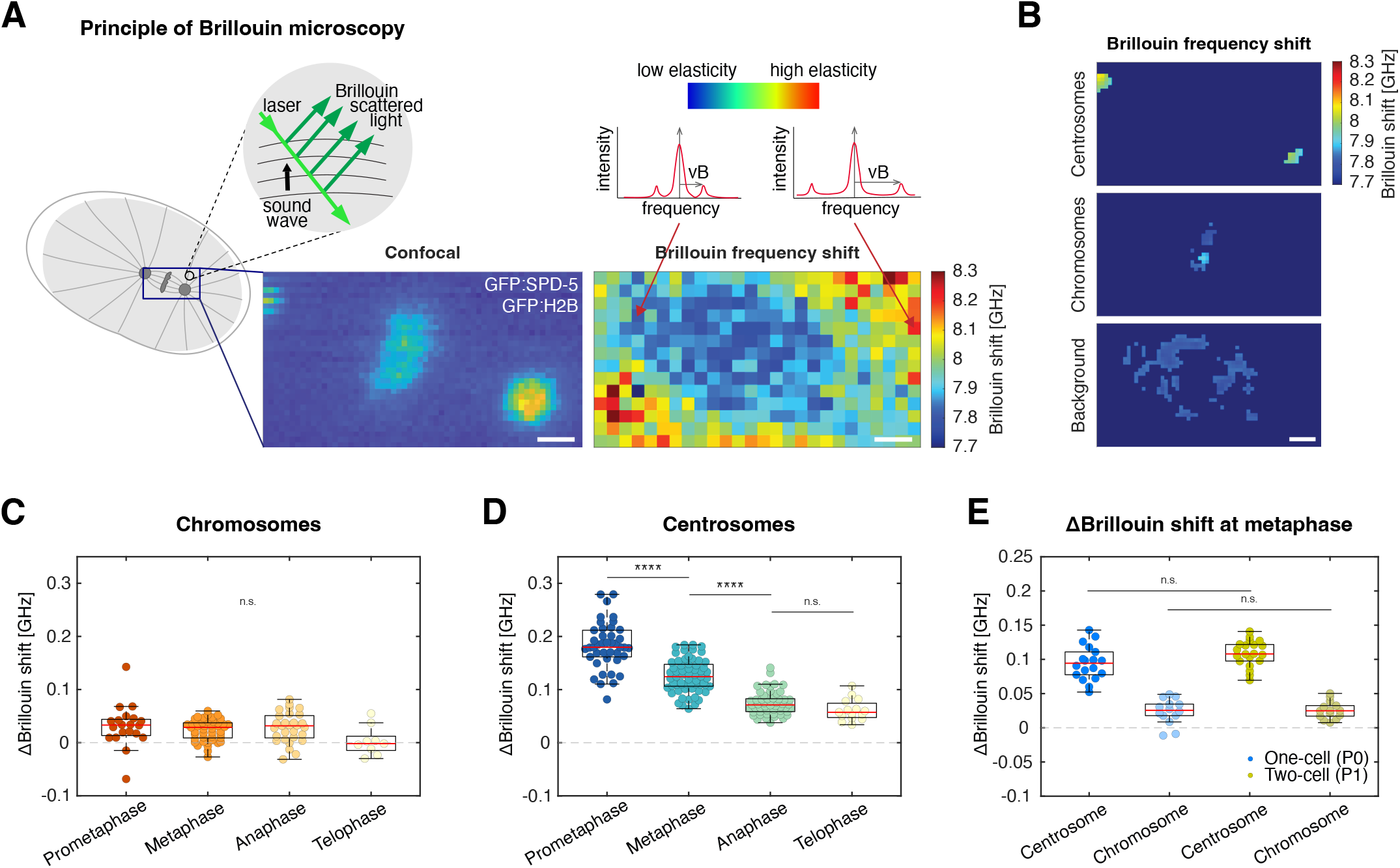
Brillouin microscopy shows softening of mitotic centrosomes *in vivo*. (A) Principle of Brillouin light scattering microscopy and application to *C. elegans* embryos. One-cell embryos were imaged using both Brillouin microscopy and confocal fluorescence microscopy modalities (GFP:SPD-5 marking centrosomes, GFP:histone H2B marking chromosomes). Scale bars: 2 µm. (B) Brillouin frequency shift for centrosome, chromosome and spindle background regions identified using fluorescence confocal signals before and after Brillouin scan in the metaphase-stage embryo shown in A. Scale bar: 2 µm. (C) Relative Brillouin shift (over background) of chromosomes remains constant throughout mitosis until decondensation at mitotic exit. (prometaphase: n = 20; metaphase: n = 45; anaphase: n = 29; telophase: n = 9; one-way ANOVA, p = 0.0672). (D) Relative Brillouin shift of centrosomes declines throughout mitosis, indicating progressive softening as early as prometaphase. (prometaphase: n = 42; metaphase: n = 63; anaphase: n = 53; telophase: n = 15; Kruskal-Wallis, p < 0.0001; post-hoc Bonferroni: prometaphase-metaphase p = 1.31e−04; metaphase-anaphase p = 4.68e−09; anaphase-telophase p = 1.00). (E) Relative Brillouin shift for centrosomes at metaphase remains constant between one and two-cell stage embryos, despite a two-fold centrosome size difference (one-cell: n =18; two-cell: n = 20; ANOVA/Tukey-Kramer post-hoc, p = 0.128). Chromosomes also show no significant change (one-cell: n = 14; two-cell: n = 14; p = 0.996). In C-E, boxplots show median (red line), IQR or interquartile range (box), and range (whiskers). Colored points depict the mean per region of interest.

Although the metaphase spindle is relatively stable, early embryos are highly dynamic and mitosis progresses rapidly. Given that our confocal Brillouin microscope requires a 100 ms effective pixel time resulting in tens of seconds to generate a full Brillouin shift map, sample movement could compromise spatial accuracy. We therefore acquired confocal fluorescence reference images before and after each Brillouin scan to confirm that no significant displacement had occurred. We then calculated the mask overlap between segmented centrosome and chromosome regions from both confocal images and spatially aligned our Brillouin measurements to these positions (Fig. 5B). Since centrosomes and chromosomes are embedded in a spindle space isolated from the surrounding embryo cytoplasm (75), we used k-means clustering to determine the characteristic shift value of the nucleoplasm/spindle space and used this as our background reference. From our measurements, both centrosomes and chromosomes displayed higher shifts (indicative of greater elasticity or higher stiffness) relative to the spindle background, with centrosomes appearing stiffer than metaphase-condensed chromosomes (Fig. S5D,E). These consistent differences demonstrate that centrosomes and chromosomes can be distinguished from their surroundings based on their distinct mechanical properties.

Having established our ability to reliably identify centrosomes and chromosomes at metaphase, we next asked whether centrosome stiffness changes during mitosis, measuring relative Brillouin shifts for both centrosomes and chromosomes from prometaphase through telophase. While chromosomes maintained constant Brillouin shift values while condensed (Fig. 5C), centrosomes showed progressive softening – exhibiting a significant decrease, not only during the known disassembly phase (anaphase to telophase; (45)), but beginning as early as prometaphase/metaphase (Fig. 5D) while the centrosome is still undergoing mitotic growth (see Fig. 1B,C). To confirm that this decrease reflects changes in mechanical properties rather than shifts in the cytoplasm-spindle boundary, we used confocal images taken immediately before and after Brillouin scans (Fig. S5FG) to reconstruct centrosome trajectories and determine their precise positions during Brillouin scan acquisition. This analysis revealed a clear relative Brillouin shift peak precisely at the predicted centrosome center across all mitotic stages (Fig. S5H), contrasting with the gradient expected from boundary transitions. This indicates that we measured centrosome-specific elasticity, not artifacts from adjacent regions. While our measurements could not resolve potential stiffness differences in prophase/S-phase centrosomes due to the inherent spatial and temporal resolution limitations of Brillouin microscopy, these results indicate centrosome softening initiates well before PCM disassembly at the end of mitosis.

To investigate whether changes in mass density during mitosis could account for the observed decrease in centrosome Brillouin shifts we performed correlative holotomography and fluorescence imaging to obtain 3D refractive index maps while localizing centrosomes and chromosomes. Since refractive index is proportional to mass density in most biological materials (76) (with lipids being an important exception (68)), these measurements provide direct quantification of mass density distribution. To validate our approach, we first measured interphase embryos at one-, two- and four-cell stages expressing GFP:histone H2B. We reproduced established density relationships (77), with nucleoplasmic density being lower than cytoplasmic density (Fig. S6A,B), confirming the reliability of our measurements. We next measured refractive index values for centrosomes, chromosomes and spindle space from prometaphase to telophase and calculated the longitudinal modulus of elasticity for each component (Fig. S6C-E). Measurement of the spindle space showed a significant increase in refractive index (Fig. S6D) and consequently longitudinal modulus (Fig. S6E) from prometaphase to telophase, potentially as a result of progressive disassembly of the spindle envelope. Both centrosomes and chromosomes exist as porous polymers embedded within a stiffening microenvironment. With their Brillouin measurements representing a composite of both centrosome/chromosome and cytoplasm, it was therefore no surprise that chromosomes followed the same trend (Fig. S6E). However, there was no similar increase for centrosomes, despite stiffening of the spindle background, confirming that condensed chromosomes maintain their stiffness while centrosomes soften progressively.

Overall, these results show that centrosomes progressively soften during mitosis, in line with their increasing deformation (Fig. 1). We next wondered whether this mechanical change correlated with their concurrent size increase. Both AFM of isolated centrosomes *in vitro* (Fig. 4H) and *in vivo* Brillouin microscopy (Fig. S6F,G) revealed a correlation between centrosome size and stiffness, with larger, mitotic, centrosomes being softer by both measures. To distinguish whether this reflected a simple size dependence or cell-cycle stage-dependent differences, we compared metaphase centrosomes from one-cell and two-cell embryos, which are two-fold different in size but at the same cell-cycle stage. Strikingly, their normalized Brillouin shifts showed no statistically significant difference (Fig. 5E). This shows that centrosome stiffness depends on cell cycle state rather than physical size alone.

Taken together, our findings demonstrate that centrosome softening is an intrinsic, cell-cycle-regulated process that initiates during early mitosis and is independent of centrosome size.

### Soft Mitotic Centrosomes May Act As Low-Pass Filters To Dampen High-Frequency Force Fluctuations

Having established that centrosome softening depends on mitotic progression rather than microtubule number or centrosome size, we next investigated its functional consequences for spindle dynamics. To explore how centrosome stiffness affects chromosome positioning and spindle length under spindle fluctuations, we developed a minimal one-dimensional model of the spindle as a spring-mass chain (Fig. 6A).

**Figure 6.**
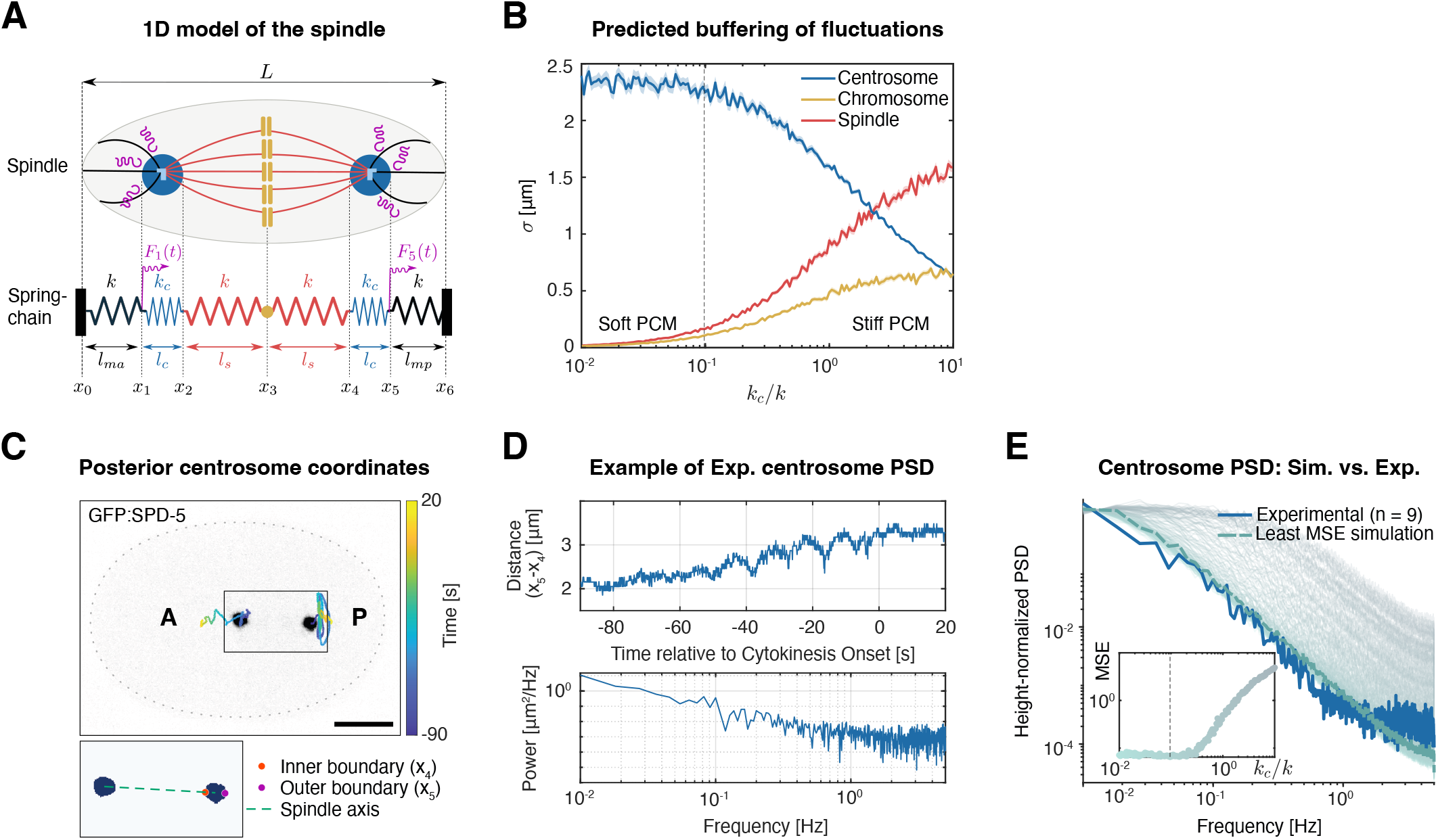
A soft PCM can filter high-frequency spindle length fluctuations. (A) Minimal 1D spring-chain model. Centrosomes are represented as PCM springs (*k*_*c*_, blue), tethered to the cortex by effective anchor springs (*k*, black) and linked to the chromosomes (yellow) by effective bridge springs (*k*, red). Random forces *F*_1_(*t*), *F*_5_(*t*) act at the outer centrosome edges (*x*_1_, *x*_5_). (B) Fluctuation amplitude versus PCM stiffness. Simulated root-mean-square (RMS) amplitudes of spindle length (σ(*x*_24_), red), centrosome length (σ(*x*_21_), blue), and chromosome midpoint (σ(*x*_3_), yellow) are plotted as a function of stiffness ratio *k*_*c*_/*k* (log-scale). In the soft-PCM regime, spindle and chromosome fluctuations are minimized, while centrosome fluctuations remain largely unchanged. Conversely, in the stiff-PCM regime, spindle and chromosome fluctuations increase substantially, and centrosome size fluctuations decrease. The vertical dashed line marks the best-fit stiffness ratio based on panel E. (C, D) Steps in power spectral density (PSD) analysis for an example one-cell stage embryo. (C) An embryo with centrosomes labeled with GFP:SPD-5 at anaphase (signal inverted for presentation). Colored lines trace the position of the two spindle poles during spindle oscillations (times relative to cytokinesis onset), highlighting the stronger oscillations seen for the posterior centrosome. Scale bar: 10 µm. Below: Segmented centrosomes indicating the inner (*x*_4_, as in A) and outer (*x*_5_, as in A) boundary points of the posterior centrosome. (D) Top: Posterior centrosome size fluctuations (calculated as the distance between points *x*_4_ and *x*_5_) over time during anaphase spindle oscillations (same embryo and time window as in C). Bottom: PSD analysis of posterior centrosome size dynamics for the same embryo. (E) Height-normalized centrosome power spectra. The experimental centrosome PSD (dark blue) is overlaid with simulation spectra spanning soft (light blue) to stiff (grey) PCM stiffness. The soft-PCM regime provides the closest match, quantified by the minimal mean-squared error (MSE) shown in the inset (log scale). The simulated PSD at this optimal value (*k*_*c*_ ≈ 0.1*k*) is highlighted by the dashed line. Stiffer PCM values increasingly over-amplify high-frequency power.

In this model, the spindle is represented by five mobile nodes (*x*_1_ through *x*_5_), bounded by fixed cortex positions at *x*_0_ = 0 (anterior) and *x*_6_ = *L* (posterior). Centrosomes are modeled as compliant springs of stiffness *k*_*c*_ and rest length *l*_*c*_, located between nodes *x*_1_ − *x*_2_ (anterior centrosome) and *x*_4_ − *x*_5_ (posterior). Each centrosome connects from its outer side (*x*_1_ or *x*_5_) to the cortex via fixed anterior (*k, l*_*ma*_) and posterior (*k, l*_*mp*_) cortical springs, and to the chromosomes at node *x*_3_ via central spindle springs (*k, l*_*s*_) (Fig. 6A). To isolate the role of PCM compliance, we set the spring constants for both the cortical and central spindle springs equal to a single value, *k*, making the centrosome stiffness *k*_*c*_ the only free mechanical parameter. We note, however, that allowing the cortical and the central spindle stiffnesses to differ would not alter the qualitative trends we observed. To test whether a compliant PCM can buffer rapid force fluctuations entering at the cortex-PCM interfaces (e.g., through astral microtubules) before they reach the central spindle and the chromosomes, we applied independent stochastic forces *F*_1_(*t*) and *F*_5_(*t*) to the outer centrosome boundaries (*x*_1_ and *x*_5_) to represent cytoplasmic fluctuations.

We simulated the overdamped dynamics of the six-spring chain (Eq. 1.1, Table S1) with centrosome stiffness *kc* spanning from 0.01*k* to 10*k*, and for simplicity assumed equal drag coefficients for all nodes (see Methods). For each distinct centrosome stiffness *k*_*c*_ value, we calculated the temporal standard deviation (simulated root-mean-square, RMS) of three key observables: centrosome length *σ*(*x*_21_), spindle length *σ*(*x*_42_) and chromosome position *σ*(*x*_3_), averaging these values across multiple simulations. Our simulations revealed that when PCM stiffness fell within 0.01 < *k*_*c*_/*k* < 0.1, spindle-length fluctuations were substantially dampened and chromosome positional variance was reduced, while centrosome length variance remained relatively unchanged (Fig. 6B). Conversely, stiffer centrosomes (*k*_*c*_/*k* ≳ 0.1) exhibited the opposite behavior: they deformed minimally (low centrosome size variance) but transmitted more force to the spindle and chromosomes, leading to increased positional variance (Fig. 6B), which could potentially destabilize kinetochore-microtubule attachments. These results suggest that soft centrosomes stabilize spindle architecture and chromosome segregation in response to force fluctuations.

Given that our theoretical work suggested that soft centrosomes could improve spindle stability and function, we next asked whether their observed *in vivo* behavior supported this soft PCM stiffness regime (Fig. 6B). To test this, we recorded centrosome dynamics during anaphase spindle oscillations (-90 to +20 s relative to cytokinesis onset) at 10 Hz in live *C. elegans* embryos expressing endogenously tagged GFP:SPD-5. We focused our analysis on the posterior centrosome, which undergoes stronger oscillations (Fig. 6C; (78)), segmenting its shape to identify the boundary points intersecting with the spindle axis (defining positions *x*_4_ and *x*_5_, Fig. 6C). By measuring temporal fluctuations in posterior centrosome size (*x*_45_(*t*) = *x*_5_(*t*) − *x*_4_(*t*)), we quantified their dynamic response through power spectral densities (PSDs), thereby mapping how centrosome shape fluctuations distribute across frequencies (Fig. 6C). Comparing simulated PSDs for different *k*_*c*_ values to this experimental data revealed the best match (smallest mean-square error, MSE) for *k*_*c*_ ≈ 0.1 *k*, in the soft PCM regime (Fig. 6B,D dashed lines). Thus, mitotic centrosomes *in vivo* exhibit soft PCM-like mechanical behavior, enabling them to filter high-frequency forces. It is important to note that in our model we applied rapid fluctuations at the cortex-PCM interfaces, which are the primary sites of astral-cortex engagement during mitosis, and intentionally restricted our analysis to this boundary-driven regime. All deformation effects and the filtering reported here occur on timescales of seconds, timescales at which the PCM behaves effectively as an elastic solid. Viscoelastic remodeling, associated with centrosome growth as characterized by Paulin et al. (51) operates on timescales well beyond our frequency band and would not alter the reported trends, although it may provide a means of additional damping of lower-frequency fluctuations.

Together, these results suggest that PCM softening during mitosis can serve as a mechanical adaptation to shield chromosomes and the central spindle from high-frequency spindle noise that could disrupt kinetochore-microtubule interactions, while transmitting sustained forces necessary for productive chromosome movements.

## Discussion

Centrosomes must adapt their size and function to meet cellular requirements, including the need to generate distinct interphase and mitotic microtubule networks while accommodating large cell volume changes during successive embryonic divisions. Here, we investigated how centrosomes respond to microtubule-generated forces and demonstrate that mitotic centrosome size scaling is controlled by mechanical deformation imposed by microtubule growth within the PCM. Our findings further reveal that mitotic deformation is not merely a consequence of increased microtubule numbers but reflects intrinsic centrosome softening, suggesting a mechanical adaptation to meet mitotic requirements (Fig. 7A).

**Figure 7.**
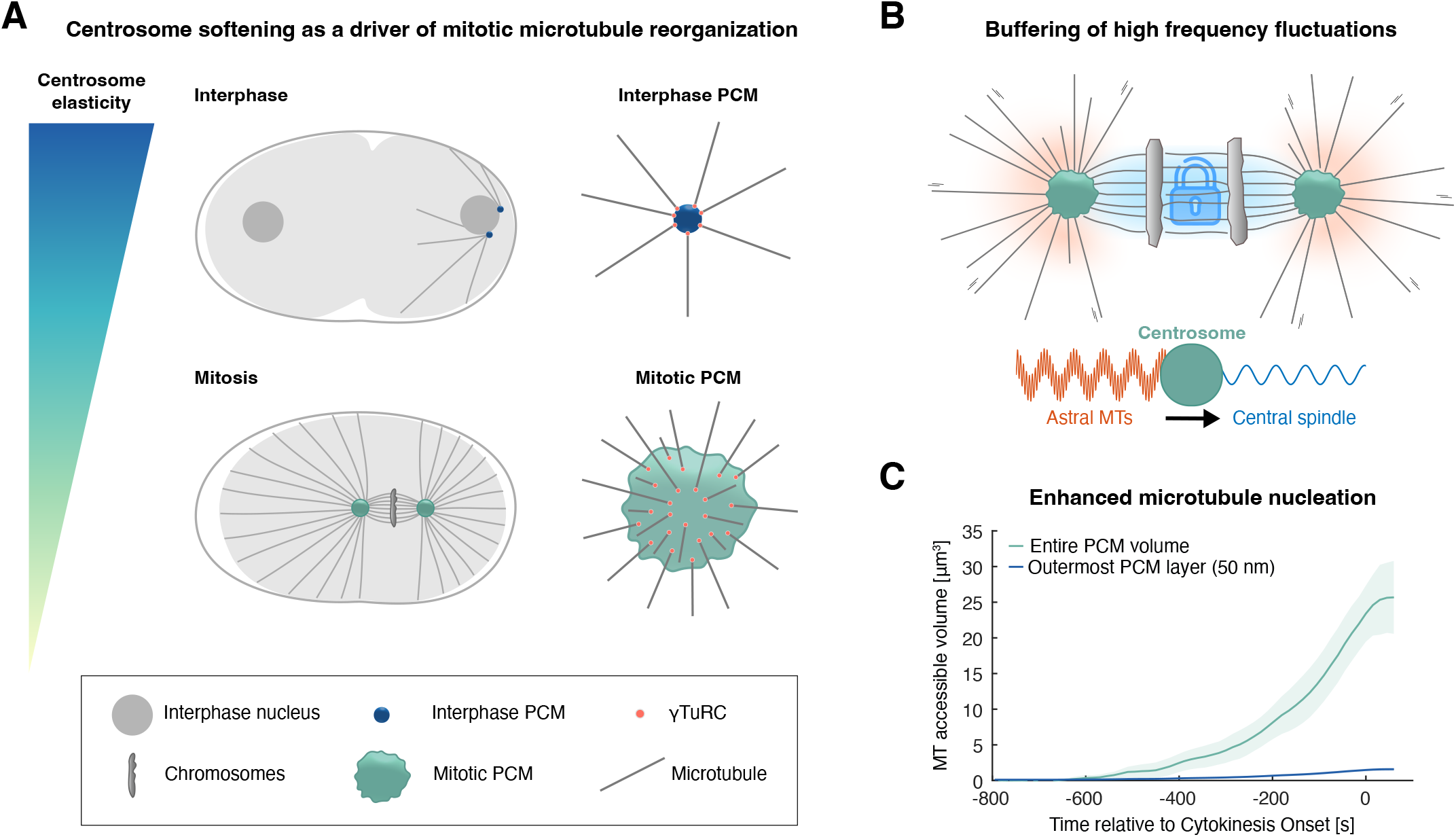
Centrosome softening as a mechanical adaptation for mitosis. (A) Summary of our findings. Mitotic centrosomes are less stiff than their interphase counterparts in the early *C. elegans* embryo. This mechanical softening correlates with a change in microtubule nucleation: microtubules emanate from the outermost PCM layer during interphase, while during mitosis, as the PCM softens, nucleation can take place deeper within the PCM (44). (B) A softer PCM could act as a low-pass filter, selectively absorbing high-frequency force fluctuations while transmitting sustained forces necessary for productive chromosome movements, thereby enabling both precise spindle positioning and proper chromosome segregation. (C) Softening could also enhance centrosomal microtubule nucleation by allowing microtubules to emanate from a larger PCM volume. If microtubules were restricted to the 50-nm-deep outer PCM layer as they are in interphase, the accessible microtubule nucleation volume would be insufficient to support spindle formation, even accounting for PCM expansion.

### Centrosome Deformation And Size Scaling During Mitosis

The early embryonic divisions of many species are rapid and characterized by minimal cell growth, leading to considerable cell volume changes as the one-cell embryo divides into smaller and smaller blastomeres. A key question is how centrosomes adapt to these changes and regulate their size accordingly. In *C. elegans* embryogenesis, centrosome size during mitosis scales proportionally with cell volume across early cleavage divisions (14), a coordination critical for proper spindle architecture and chromosome segregation (16). While previous work proposed that centrosome size is constrained by limiting amounts of centrosomal material (with SPD-2 as the limiting factor (14)), the absence of scaling in nocodazole-treated centrosomes during the first embryonic divisions argues against this model, as scaffold polymer amounts remain near-constant across multiple divisions rather than halving each time (Fig. S1). Our data instead support a microtubule-dependent mechanism. We show that forces from growing microtubules deform the PCM scaffold from within, with the resulting deformation scaling with cell size (Figs. 1,3).

In our revised model centrosome size is instead determined by microtubule numbers, which appear to scale with cell volume. This scaling relationship emerges because the cytoplasmic pool of available tubulin and microtubule regulators, while maintained at constant levels prior to zygotic genome activation, shrinks as cell volume decreases (18, 19). Across diverse systems, microtubule dynamics are key determinants of spindle length scaling: in *C. elegans* (63), *Paracentrotus lividus* (63) and *Xenopus laevis* (79, 80), reduced spindle lengths correlate with declining microtubule growth rates, likely mediated by limiting amounts of microtubule regulators like XMAP215 (81). Conversely, increasing levels of purified tubulin in *Xenopus* egg extracts led to an increase in spindle length and mass (82), implicating tubulin limitation as a scaling driver. This aligns with the established dependence of microtubule polymerization rates on free tubulin concentration *in vitro* (83, 84). Thus, whether through tubulin availability or other microtubule regulatory factors, microtubule-dependent scaling helps explain the observed coordination between spindle and centrosome size (16), which emerges from their mutual dependence on microtubule numbers and aligns with the role of centrosomal factors in controlling spindle assembly (16, 85).

Why can cortical pulling forces affect centrosome shape but not override its size regulation (Fig. 2)? We hypothesize that this reflects differences in the force generation capacity and spatial distribution between internal and external forces. Within the PCM, individual microtubules can generate up to 3-4 pN of pushing force through polymerization (86-88), with shorter microtubules – like those found within the PCM (7) being particularly effective due to their higher resistance to buckling (86). With over 5,000 microtubules per metaphase centrosome in *C. elegans* one-cell stage embryos (89), the collective pushing forces can be substantial. In contrast, cortical pulling forces are comparatively weaker despite each cortical dynein complex generating up to 5 pN of force via dynein-mediated depolymerization (90-92), because only ∼200 microtubules contact the cortex at any given time (93). However, since cortical force generators are asymmetrically positioned within the embryo (38), they can impact centrosome shape, whereas astral microtubules radiate uniformly outward and thus impose no directional bias on centrosome deformation (Fig. S2).

While our work establishes this microtubule-dependent mechanical scaling model in *C. elegans* embryos, whether this represents a conserved metazoan strategy remains to be tested. The elegance of this mechanism lies in its simplicity – by modulating microtubule numbers rather than centrosome assembly, cells can rapidly coordinate spindle and centrosome dimensions with cell volume. Our findings thus reveal a potentially universal biophysical solution for centrosome and spindle size scaling.

### Centrosome Softening During Mitosis

While decades of research have shed light on the centrosome’s molecular composition – with hundreds of components regulating its biochemical functions – its mechanical behavior under physiological forces has remained largely unexplored. This knowledge gap is particularly striking given the centrosome’s role as the mechanical anchor for the spindle, a force-generating machine that exerts hundreds of piconewtons during chromosome segregation. These high mechanical demands make centrosome material properties crucial for successful division. The study of centrosome mechanics has lagged behind its molecular characterization due to the technical challenge of probing intracellular mechanics without disrupting cell division. This limitation has left us with an incomplete picture of how the PCM achieves dynamic growth, yet resists microtubule forces throughout the cell cycle. As predicted by Paulin et al. (51), a PCM with insufficient elasticity would be prone to mechanical failure, while one that is too stiff would resist the expansion necessary for mitotic growth. Prior work using shear stress induced by optically generated flows showed that centrosomes mechanically weaken during their disassembly phase, but the material properties of mature metaphase centrosomes could not be assessed (45). Thus, it remained unknown whether centrosomes *in vivo* transition from a liquid-like to a solid state, as suggested by studies on *in vitro* SPD-5 condensates (26) or undergo alternative mechanical transitions. Here, using techniques not previously applied to centrosomes, AFM and Brillouin microscopy, we demonstrate that centrosomes undergo a mechanical transition during mitosis, becoming progressively softer (Fig. 5). However, and in contrast to *in vitro* predictions, both interphase and mitotic centrosomes exhibit solid-like properties (Fig. 4), enabling them to resist microtubule forces.

While AFM is the gold standard in quantitative mechanobiology, it cannot be used to probe intracellular structures such as centrosomes inside the living cell. Unlike AFM, Brillouin microscopy is not surface-limited and therefore holds great promise for use within cells *in vivo* at near diffraction-limited resolution. However, the application of Brillouin microscopy to biological systems is not without its limitations. The Brillouin shift depends on both refractive index and mass density, not just mechanics; although we took all possible precautions – measuring refractive index at near-confocal resolution to approximate density and extract the longitudinal modulus – a key limitation is that both parameters were derived via the same quantitative phase imaging method. Furthermore, relating the Young’s modulus obtained by AFM to the Brillouin longitudinal modulus is not straightforward. The two techniques probe different time- and length-scales and their moduli are mathematically comparable only for linear isotropic materials, unlike the complex nature of biological materials (94). It is worth noting, however, that mitotic centrosome softening was observed with both techniques. As one of the few studies to apply both modalities to the same biological question, this strengthens our conclusions and underscores the value of AFM and Brillouin microscopy as complementary tools.

Why do centrosomes soften specifically during mitosis? This cell cycle-regulated mechanical transition appears counterintuitive given the centrosome’s role as the spindle’s anchor point. However, our data suggest mitotic softening serves a critical function: it converts the centrosome into a biological low-pass filter (Fig. 6). By selectively absorbing high-frequency force fluctuations (e.g., from microtubule dynamics or cortical pulling) while permitting slow, controlled movements, this soft state may enable both precise spindle positioning and proper chromosome segregation (Fig. 7B). This filtering capacity may be a crucial requirement for achieving the remarkable precision of spindle centering in *C. elegans* embryos (78). Our theoretical model shows that excessively stiff centrosomes would lose this buffering ability, potentially impacting chromosome positioning and spindle length regulation (Fig. 6). Mitotic chromosome movement may be aided by another feature, their relative isolation from the general cytoplasm. Using Brillouin microscopy and quantitative phase imaging to assess refractive index, we found that the spindle space in *C. elegans* embryos is less dense than the surrounding cytoplasm (Fig. 5). While animals undergo open mitosis, they retain envelope-like membranes surrounding the mitotic spindle (95-97) – a feature observed not only in embryonic systems (e.g., *Drosophila* (74), *C. elegans* (75), sea urchins (73, 98), but also in somatic cells (99). Given the extreme crowding of the embryonic cytoplasm (packed with maternally inherited material), this partitioning may be particularly critical during embryogenesis, as it creates a less crowded environment to ensure unobstructed chromosome segregation. Future studies should further characterize the functional significance of this cytoplasmic compartmentalization.

If there are benefits to soft centrosomes in mitosis, why would centrosomes not remain equally soft throughout the entire cell cycle? Stiffer interphase centrosomes may offer distinct advantages, including providing structural resilience against deformation in the crowded cytoplasm, enabling more effective anchoring of interphase microtubules, and facilitating more efficient force transmission for rapid organelle positioning and the transmission of mechanical signals to the nucleus. This stiffness regulation suggests that centrosomes dynamically adapt their material properties to cell cycle stage-specific demands – softer during mitosis to function as a protective mechanical filter and stiffer during interphase to maintain structural integrity and facilitate force transmission.

Finally, another functional interpretation of mitotic centrosome softening relates to microtubule organization. In interphase, microtubule nucleation occurs primarily in the outermost PCM layer (44), whereas during mitosis microtubules permeate most of the PCM volume (6, 7, 44). This dramatic reorganization raises important questions about how PCM mechanical properties regulate microtubule nucleation capacity. Our findings that centrosomes soften during mitosis suggest that this mechanical transition may facilitate microtubule nucleation within an expanded PCM scaffold. Specifically, a softer, more flexible PCM could better accommodate and sustain the polymerization of increased microtubule numbers – a fundamental requirement for proper spindle assembly. Critically, if microtubules were restricted to the 50-nm-deep outer PCM layer (as in interphase; (44)), even accounting for mitotic PCM expansion, the accessible microtubule nucleation volume would be insufficient to support spindle formation (Fig. 7C).

While our work has revealed that centrosomes undergo a cell-cycle regulated transition to a softer state during mitosis, an important open question remains regarding the molecular mechanisms controlling this process. While several regulators of PCM scaffold assembly and integrity have been identified, notably PCMD-1, PLK-1 and SPD-2, their inhibition typically weakens the centrosome structure. For example, PCMD-1 – a critical linker between PCM and centrioles – maintains PCM integrity against microtubule-dependent forces (100-102), in line with the centriole’s structural role. Similarly, partial PLK-1 inhibition or a temperature-sensitive SPD-2 mutation rendered centrosomes susceptible to deformation via ectopic cytoplasmic flows during metaphase, whereas control centrosomes remained unaffected (45). Consistent with this stabilizing role, both proteins depart from the PCM during anaphase, preceding disassembly (45). The recent discovery that PLK-1-mediated phosphorylation of SPD-5 changes the viscoelasticity of mitotic PCM towards more solid-like properties (103) suggests a potential antagonistic relationship with the softening transition. Future studies should determine whether PLK-1 phosphorylation directly opposes centrosome softening or operates through independent pathways.

In conclusion, our work redefines centrosomes as dynamic, mechanically active organelles that tune their properties to meet cell stage-specific demands, moving beyond their traditional view as static structures or simple biomolecular hubs. These findings open up new questions about the specific molecular mechanisms that drive centrosome mitotic softening and how its dysregulation might contribute to errors in chromosome segregation.

## Materials and Methods

### *C. elegans* Strains and Culture Conditions

The following strains were used in this study:

AMJ795: *rme-2(b1008) IV; pwIs23[Pvit-2::vit-2::gfp]* (104)

BCN9071: *vit-2(crg9070[vit-2::gfp]) X* (105)

DAM946: *spd-5(vie26[gfp::spd-5 +loxP]) I; unc-119(ed3) III; ltIs37[pAA64; Ppie-1::mcherry::his-58; unc-119(+)] IV; ltIs69[pAA191; Ppie-1::mcherry::spd-2; unc-119(+)] IV* (34)

DAM1075: *spd-5(vie26[gfp::spd-5 +loxP]) I* (106)

DAM1342: *spd-5(vie26[gfp::spd-5 +loxP]) I* ; *unc-119(ed3) III; ruIs32[pAZ132; Ppie-1::gfp::his-58] III* (This study)

DH1390: *rme-2(b1008) IV* (70)

N2: wild-type

SA854: *tbb-2(tj26[gfp::tbb-2]) III* (107)

XA3501: *unc-119(ed3) III; ruIs32[pAZ132; Ppie-1::gfp::his-58] III; ruIs57[pAZ147; Ppie-1::gfp::tbb-2] V* (108)

All strains were maintained at 20 °C.

### RNA-Mediated Interference

RNAi experiments were performed by soaking (109). For *gpr-1/2* and *efa-6* RNAi, standard RNAi conditions of 48 h at 16 °C were used to ensure full depletion. For *klp-7* RNAi, recovery time was slightly reduced to 24 h at 20 °C with an additional 18 h recovery at 16 °C. For eggshell permeabilization by *perm-1* RNAi in combination with *mdf-2* depletion, recovery time was reduced to 24 h at 20 °C to minimize phenotypes beyond embryo permeability.

dsRNAs used were as follows:

*efa-6*: 2.6 mg/ml, amplified from N2 genomic DNA

Forward primer: AATTAACCCTCACTAAAGGGGACACTCCGTCGAAACATT

Reverse primer: TAATACGACTCACTATAGGCCGTCTTGATGTTGAAGCAA

*gpr-1/2*: 3.9 mg/ml, amplified from N2 genomic DNA

Forward primer: TAATACGACTCACTATAGGAGCATGTGATTCCACACGTC

Reverse primer: AATTAACCCTCACTAAAGGTCTGGCAGCAGACAGTTCAG

*klp-7*: 1.6 mg/ml, amplified from N2 genomic DNA

Forward primer: AATTAACCCTCACTAAAGGGTCAACGCGTAGATTCCCAT

Reverse primer: TAATACGACTCACTATAGGCCCGCCACTGACTAATGTTT

*mdf-2*: 1.7 mg/ml, amplified from N2 genomic DNA

Forward primer: TAATACGACTCACTATAGGCAGATTTCGGCCTTAACAGC

Reverse primer: AATTAACCCTCACTAAAGGCTTTTTGAGGGGGTTTGTGA

*perm-1*: 3.2 mg/ml, amplified from N2 genomic DNA

Forward primer: AATTAACCCTCACTAAAGGAATTTTCTAGGTCGTCAATCTTCA

Reverse primer: TAATACGACTCACTATAGGCGAAAACGCGATCATTTTTA

### Centrosome Isolations from *C. elegans* Embryos

Worms expressing GFP:SPD-5 (DAM1075) were grown in synchronous liquid culture (110) to young adult stage (5-10 embryos/ worm) and bleached to harvest embryos. Embryos were washed 3x with cold M9 before drop freezing in liquid nitrogen. 2 g of frozen embryos were thawed in 14 mL of HB buffer (20 mM HEPES, pH 7.8; 5 mM K-acetate; 0.5 mM MgCl_2_; 0.5 mM DTT) supplemented with protease inhibitors cOmplete Mini EDTA-free Protease Inhibitor Cocktail (Roche, Cat# 11836170001, 1 tablet/10 ml lysis buffer), 1 mM PMSF, 1 mM Benzodiazepine, and 0.01x PhosSTOP (Roche, Cat# 4906845001) and subjected to tip sonication (3 × 15 s at 30% continuous output, 3 × 15 s at 40% output, Bandelin Sonopuls GM70) with 30 s cooling on ice between cycles. Lysates were filtered 2x through Miracloth (Merck, Cat# 475855) and clarified by centrifugation 2x at 1,500 x g for 10 min at 4 °C. 10 ml of cleared supernatant was mixed with 28 ml 70% sucrose to bring the sucrose concentration to 45%, layered onto a sucrose step gradient (3 ml 70% sucrose, 4 ml 55% sucrose, both containing 1 mM GTP) in thin-wall ultracentrifuge tubes (Beckman Coulter, Cat# 326823) and centrifuged at 100,000 x g for 2 h at 4 °C in a swinging bucket rotor (SW28, Beckman Coulter), using slow acceleration and deceleration. Fractions were collected from the bottom of the tube using a 25G needle in 500 µl fractions. Centrosomes were counted in each fraction by fluorescence microscopy using a Neubauer counting chamber and total protein concentration was determined using the Bradford assay.

### *In Vitro* Microtubule Nucleation Experiments

Centrosome integrity was confirmed by assessing their nucleation capacity. For this, porcine brain tubulin (52 mg/ml) was diluted 1:5 in cold tubulin dilution buffer TDB (20 mM K-PIPES, pH 6.8; 1 mM MgCl_2_; 2 mM EGTA; 10% glycerol; 1 mM GTP) to a final concentration of 10 mg/ml and 10 mg/ml rhodamine-labeled porcine tubulin (Cytoskeleton, Cat# TL590M) was added at a ratio of 1:20 to prepare a red-tubulin mix. Microtubule nucleation assays were carried out in custom flow chambers assembled using double-sided tape and acid-treated coverslips (see Fig. 4C). Chambers were blocked for 10 min with 1% BSA in TDB. Centrosomes diluted 1:1 with BRB80 (20 mM K-PIPES, pH 6.8; 1 mM MgCl_2_; 2 mM EGTA) were introduced and allowed to adhere for 10 min. Chambers were then washed with TDB before incubating with 10 µl of red-tubulin mix in a humidified chamber at 29 °C for 10 min. Reactions were stopped by introducing 10 µl of fixation buffer (1% glutaraldehyde in BRB80) pre-warmed to 29 °C and incubating for 3 min at room temperature. Finally, chambers were washed with TDB, mounted in Vectashield and sealed with nail polish for imaging.

Slides were examined using a Zeiss Axio Imager Z2 microscope equipped with an 40x 1.3 NA EC Plan-Neofluar oil immersion objective and a Lumencor SOLA SE II light source. Fluorescent signals were detected using Filter Sets 38 for GFP (excitation BP 470/40, beamsplitter FT 495, emission BP 525/50) and 43 for Rhodamine (excitation BP 545/25, beamsplitter FT 570, emission BP 605/70) and GFP/Rhodamine image stacks were acquired using a Photometrics CoolSNAP-HQ2 cooled CCD camera controlled by Zeiss ZEN 2 blue software.

### Spinning Disk Confocal Microscopy

Live cell imaging of centrosome dynamics was performed using a Yokogawa CSU-X1 spinning disk confocal mounted on a Zeiss Axio Observer Z1 microscope equipped with 100 mW 488 nm and 561 nm solid-state lasers and an Excelitas pco.edge 4.2 sCMOS camera and controlled by VisiView 6.0 software (Visitron Systems). For the imaging in Figs. 1, S1, S5A and 6, we used a 63x 1.4 NA Plan-Apochromat oil immersion objective with DIC III optics. For Figs. 2, S2, 3 and S3, data were acquired using a 63x 1.3 NA LCI Plan-Neofluar glycerol/water immersion objective with DIC III optics and a correction collar, optimized for the refractive index of the mounting medium. Most experiments used 150 ms exposure time for the 488 nm channel, 250 ms for the 561 nm channel and 400 ms for the DIC channel. For cytoplasmic yolk imaging in Fig. S5A, exposure time for the 488 nm channel was increased to 250 ms, while for the high temporal resolution imaging in Fig. 6, exposure time was reduced to 100 ms. Laser power was kept constant across all imaging sessions and experiments.

For acute nocodazole treatments, *perm-1;mdf-2* double RNAi-treated worms co-expressing GFP:SPD-5, mCherry:SPD-2 and mCherry:Histone (DAM946) were dissected onto 24 × 60 mm coverslips in an 8 μl droplet of meiosis medium (60% Leibovitz’s L-15 medium, 25 mM HEPES pH 7.4, 0.5% inulin, 20% heat-inactivated fetal bovine serum) containing 15.7 μm polystyrene spacer beads (Polysciences, Cat # 18328-5). An 18 × 18 mm coverslip was then placed on top to immobilize embryos between the two coverslips. The sample was sealed at both ends with two-component dental silicone (Picodent eco-sil speed). 33 × 0.5 µm GFP z-stacks along with single midplane mCherry and DIC images were acquired every 15s using low laser illumination to minimize photobleaching. For drug application in S-phase (post-centrosome separation) or at NEBD, meiosis medium was replaced with medium containing 0.01 μg/μl nocodazole by pipetting the nocodazole-containing medium onto one side of the horizontal opening between the two coverslips while simultaneously drawing off the original medium from the opposite side using filter paper. Embryos requiring > 3 min for complete medium exchange were excluded from further analysis. Controls for nocodazole treatments at the one-cell stage were *perm-1;mdf-2* double RNAi embryos maintained in drug-free meiosis medium. These embryos showed no significant differences in cell cycle timing (from male pronuclear permeabilization to chromosome decondensation) compared to untreated one-cell stage embryos (n=14 RNAi embryos, 16 untreated; Mann-Whitney U test, p = 0.2436). Furthermore, analysis of centrosome dynamics using a linear mixed-effects model confirmed no significant differences in either centrosome size (p = 0.6576) or GFP:SPD-5 fluorescence intensity (p = 0.1745). Because *perm-1* RNAi induces polar body resorption that could potentially affect chromosome segregation (47), we used untreated embryos for control comparisons at the two- and four-cell stages, as the two control conditions behaved similarly at the one-cell stage.

For the experiments in Figs. 2, S2, 3 and S3, young adult animals were dissected onto 24 × 60 mm coverslips in an 8 µl droplet of 30% iodixanol (OptiPrep, STEMCELL Technologies, Cat # 07820) in M9 (refractive index matched to embryos, n = 1.365) containing 24.7 μm polystyrene spacer beads (Polysciences, Cat # 07313-5) to minimize spherical aberration during imaging. Samples were mounted under an 18 × 18 mm coverslip and sealed with silicone. For embryos co-expressing GFP:SPD-5, mCherry:SPD-2 and mCherry:histone H2B (DAM946), 33 × 0.5 µm GFP z-stacks along with single midplane mCherry and DIC images were acquired every 15s using low laser illumination to minimize photobleaching. For embryos expressing GFP:Tubulin only (SA854), the same imaging conditions were used except leaving out the mCherry channel. For embryos co-expressing GFP:Tubulin and GFP:histone H2B (XA3501), 17 × 0.5 µm GFP z-stacks were acquired along with a DIC image at the midplane.

For yolk imaging in Fig. S5A, dissected embryos were filmed in an 8 μl droplet of meiosis medium (60% Leibovitz’s L-15 medium, 25 mM HEPES pH 7.4, 0.5% inulin, 20% heat-inactivated fetal bovine serum). Samples were mounted on 24 × 60 mm coverslips taped to a metal holder. To prevent compression and minimize medium evaporation, vaseline ring spacers were used to support an overlying 18 × 18 mm coverslip. GFP z-stacks were acquired every 15s over an 8 µm range using 17 steps at 0.5 µm z-spacing along with a midplane DIC image.

For the examination of centrosome dynamics at high temporal resolution (Fig. 6), young adult animals co-expressing GFP:SPD-5 and mCherry:Histone (DAM946) were dissected in M9 containing 15.7 μm polystyrene spacer beads (Polysciences, Cat # 18328-5) to restrict spindle movement in z, ensuring centrosomes remained within the plane of imaging. Single plane GFP images were acquired in continuous acquisition mode.

### Centrosome Mechanical Characterization by AFM

AFM experiments were performed using a Bruker NanoWizard ULTRA Speed A AFM system integrated with a Zeiss Axio Observer D1 fluorescence microscope used to initially locate centrosomes. For sample preparation, acid-washed coverslips (24 × 60 mm) were assembled with 3D-printed rectangular chambers using two-component dental silicone (Picodent twinduo extrahard) to seal the chamber system. Isolated centrosomes in sucrose were allowed to attach to the glass surface for 10 minutes under a small (10 mm diameter) round coverslip. The sample was then immersed in BRB80 buffer, and the coverslip removed.

Nanoindentation tests in Quantitative Imaging (QI) mode were performed using an MSCT-10D cantilever (Bruker) with a nominal spring constant of 0.03 N/m (measured at 0.027 N/m in air via thermal noise calibration (111, 112)) and a resonance frequency of 15 kHz. Force volume map measurements were conducted using tipless MLCT-O10 cantilevers (Bruker) with nominal spring constants of 0.1 N/m (measured at 0.066 N/m and 0.065 N/m in air for MLCT-O10-7E and MLCT-O10-4E, respectively) and a nominal resonance frequency of 38 kHz. These cantilevers were furnished with a 2 μm diameter borosilicate bead (Duke Standards, Thermo Fisher, Cat# 9002) using a two-component epoxy glue (UHU Plus), as previously described (66, 113). Cantilever spring constant calibration was performed in air via the thermal noise method (111, 112) prior to both mechanical characterization and furnishing the tipless cantilevers with the colloidal probes. For colloidal probes, the pre-calibrated spring constant was used. Employing established methods in the Thurner lab, tip shape characterization was performed prior to mechanical tests for both nanosized tips on the MSCT-10D cantilevers (114) and the colloidal probes furnished on the MLCT-O10 cantilevers (66). Briefly, a conical reference spike with a 50° opening angle and 5 nm tip radius from a TGT1 calibration grating (NT-MDT, Spectrum Instruments) was scanned in QI mode. For nanosized AFM tips, the resulting image represents the envelope of the TGT1 spike and the AFM tip. The AFM tip was reconstructed from the envelope image using a custom MATLAB code (114) (available at: https://github.com/Rufman91/ForceMapAnalysis) implementing a reconstruction algorithm developed by (115). For colloidal probes, as previously reported (66) the resulting envelope image represents the true geometry of the colloidal probe. The height data corresponding to the colloidal probe’s geometry were then used in the Sneddon method to calculate the indentation modulus.

Prior to centrosome measurements, cantilever deflection sensitivity was calibrated in BRB80 buffer using a contact-based approach. This involved recording the vertical deflection (in meters) as a function of z-displacement (in meters) on a glass slide. Calibration of the cantilever deflection sensitivity was performed iteratively until the sensitivity value converged. Scanning in QI mode (nanoindentation tests) was performed at 0.3 nN applied force, with 1000 nm z-length, at 200 ms per pixel, and a 13.36 nm pixel size. Force volume maps (measurements with a colloidal probe) were recorded at 1.5 nN relative applied force, with 1000 nm z-length, at 20 μm/s z-displacement speed, and 10 kHz scan rate. The pixel size was adjusted between 19.36 and 93.23 nm to match individual centrosome dimensions.

### Brillouin Microscopy

The microscope setup used for Brillouin microscopy combines point-scanning fluorescence confocal imaging with Brillouin spectroscopy, as previously described (116). In brief, a custom-built spectrometer was coupled to a Zeiss Axiovert 200/M microscope body. The spectrometer is based on a two-stage VIPA configuration (117) with the addition of a Lyot stop to increase the suppression of the elastically scattered light (118).

Imaging was performed with 5 mW laser power (measured before the objective) through a 40x oil-immersion objective (NA 1.0) with an exposure time of 120 ms per point. Brillouin scans were acquired with a 0.6 µm step size across rectangular regions of interest (ROIs) covering the mitotic spindle, typically requiring ∼40s per Brillouin scan. Confocal images (with an effective pixel size of 0.25 µm) were taken immediately before and after each Brillouin scan to confirm that no detectable displacement of centrosomes or chromosomes occurred during acquisition.

All data were acquired using a 532 nm Brillouin wavelength laser, with the exception of the one- and two-cell stage metaphase dataset in Figure 5E, which employed a 660 nm laser.

The acquisition was performed using custom LabVIEW software, which enabled ROI selection based on brightfield images and subsequent acquisition of both fluorescence confocal and Brillouin images. The Brillouin shift *νB* was calculated by performing a Lorentzian fit to the raw Brillouin spectra, using distilled water as the reference material for calibration.

For sample preparation, *C. elegans* adults co-expressing GFP:SPD-5 and GFP:Histone (DAM1342) or VIT-2:GFP in a wild-type (BCN9071) or *rme-2* mutant background (AMJ765) were dissected in 20 µl of M9 buffer and mounted in 35 mm glass-bottom dishes (1.5 mm thickness). A custom incubator maintained samples at 20 °C during imaging to minimize thermal drift and slow down cell cycle progression.

To enable reliable anaphase and telophase centrosome measurements – stages where spindle rocking disrupts confocal fluorescence image alignment and prevents centrosome and chromosome localization during the Brillouin scan – worms were treated with *gpr-1/2* RNAi to suppress spindle oscillations. This treatment did not significantly alter centrosome (ANOVA, p = 0.20317) or chromosome ΔBrillouin shifts (Kruskal-Wallis, p = 0.34956) between RNAi-treated and untreated conditions.

### Nanolive Microscopy

Refractive index (RI) tomograms and fluorescence images were acquired using a Nanolive 3D Cell Explorer Fluo microscope equipped with a DAPI-FITC-TRITC/Cy5 module. For fluorescence imaging, we used FITC filter settings with 30 gain, 60% LED intensity, and 150 ms exposure time. Adult *C. elegans* were dissected in 8 µl of M9 buffer containing 20% iodixanol (RI = 1.3659) to match the cytoplasmic refractive index of embryos (77, 119) and 15.7 μm polystyrene spacer beads (Polysciences, Cat # 18328-5) to reduce embryo thickness and minimize scattering artifacts. Embryo samples were mounted between a 24 × 60 mm bottom coverslip and an 18 × 18 mm top coverslip.

### Analysis of Spinning Disk Confocal Centrosome Image Sequences

Image stacks were processed using Fiji/ImageJ (version 2.14.0/1.54f), where channels were split and timestamp metadata extracted using the Bio-Formats Timestamp Reader plugin (available at https://www.maxperutzlabs.ac.at/research/facilities/biooptics-light-microscopy). Processed movie sequences were saved as TIFF files for MATLAB analysis (R2021b). Temporal alignment of movies was achieved by synchronizing datasets to two reference time points: 1) male pronuclear /nuclear permeabilization for S-phase-treated centrosomes, and 2) chromosome decondensation for NEBD-treated embryos. Nuclear permeabilization was defined as the frame preceding complete cytoplasmic GFP invasion of the nuclear space. Chromosome decondensation timing was determined using a modified version of the method by Maddox et al. (120). Briefly, a custom MATLAB script quantified fluorescence intensity changes within a 50 × 50 px nuclear region. Decondensation onset was defined as the first frame showing a decrease in the percentage of pixels below three intensity thresholds (80%, 85% and 90% of maximum nuclear fluorescence), indicating chromatin decompaction.

Spherical aberration correction was performed using the Fiji macro focal_shift_correction.txt (121). For oil-immersion objectives (NA = 1.4, immersion medium RI = 1.52, mounting medium RI = 1.33), a median focal shift correction factor of 0.838 x was applied. For glycerol-immersion objectives (NA = 1.3, immersion medium RI = 1.47, mounting medium RI = 1.365), a median correction factor of 0.910 x was used. For each movie, camera noise was corrected by subtracting the mean pixel intensity of out of embryo background from all frames. Finally, to account for fluorescence photobleaching in time-lapse sequences, an exponential model (*Y* = *a* · *e*^*bx*^ + *c*) was fit to a 5-frame moving average of the GFP signal intensity to correct for fluorescence decay. The model constrained amplitudes (a, c ≥ 0) and decay rates (b ≤ 0), excluding pre-corrected or short sequences (<3 frames). Correction factors were validated via R^2^ and RMSE calculations, with correction factors saved for each frame. Interactive plots allowed manual verification of both raw data and corrected intensities before proceeding.

For centrosome segmentation, centrosome coordinates were initially determined by first manually selecting the embryo region using the maximum intensity projection of the GFP:SPD-5 channel across all timepoints, followed by automatic segmentation to identify centrosome coordinates. If the algorithm detected fewer or more than two centrosomes, the user was prompted to manually correct the selection. Coordinates in the embryo cytoplasm (away from the centrosomes) were similarly defined using the maximum intensity projection across time. These centrosome coordinates were then used to define a prismatic volume (60-pixel diameter across all z-stacks) around each centrosome, reducing image complexity for intensity-based segmentation. The centrosome midplane was identified by summing intensity values within the defined volume for each z-plane, and the plane with the maximum intensity sum was selected. For segmentation, mild Gaussian filtering (kernel size: 3 × 3, σ = 1) was applied in both 2D and 3D, followed by Otsu thresholding (using MATLAB’s multithresh function) for 2D and 3D segmentation. Morphological properties including refined centroid coordinates, equivalent diameter, orientation and area/volume measurements were extracted using MATLAB’s regionprops (2D) or regionprops3 (3D) functions. Anterior and posterior centrosomes were assigned based on their relative distances to user-defined posterior coordinates.

Centrosome fluorescence intensity measurements (mean and integrated intensity) were obtained by applying 2D or 3D segmentation masks to the original z-stack images after subtracting the cytoplasmic background. The background was calculated as the mean intensity in squares sized according to developmental stage (60 × 60 px for one-cell, 42 px for two-cell, and 24 px for four-cell embryos). For closely spaced centrosomes at the 2D midplane before separation, we implemented a watershed transform algorithm (MATLAB’s watershed function) to resolve individual centrosomes. The algorithm first computed the Euclidean distance transform of the binary mask (bwdist), then applied watershed segmentation with oversegmentation correction (merging regions when more than two objects were detected). Resulting parameters from separated centrosomes were saved individually. To ensure consistency, 3D centrosome parameters were only retained when the number of detected centrosomes matched between 2D and 3D analyses, with watershed processing exclusively applied to the 2D midplane projection. We excluded centrosomes that were either truncated or out of focus by analyzing their 3D spatial distribution. Centrosomes were flagged for exclusion if they either touched the top or bottom boundaries of the z-stack or showed >20% surface loss (calculated via 26-connectivity 3D surface voxel analysis). For excluded centrosomes, all associated 2D and 3D measurements (including intensity, volume and geometric parameters) were set to NaN to prevent biased quantification.

To validate 2D segmentation accuracy at the midplane, we performed radial intensity profiling of centrosomal intensity distributions in elliptical coordinates. Using the segmented centroid position and orientation, we generated concentric elliptical rings centered on the centrosome and calculated mean intensity values within each ring. The resulting radial intensity profile was mirrored to enforce symmetry and fitted with a Gaussian function 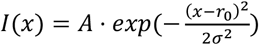. From this fit, we determined the intensity distribution’s full-width half-maximum (FWHM), which was converted to a physical radius measurement.

We evaluated centrosome radius measurements by comparing 2D intensity-based segmentation with two alternative approaches: radial intensity profiling and full 3D segmentation. For each of the 18 experimental datasets, anterior and posterior centrosome means were pooled for regression analysis (MATLAB fitlm). Comparison between 2D segmentation and radial intensity profiling revealed a strong proportional relationship: Radius [profiling] = 0.984 · Radius [segmentation] - 0.025 μm with 95% CIs of [0.978, 0.991] for the slope (1.55% deviation from 1) and [-0.031, -0.019] μm for the intercept and high R^2^ = 0.976 (Fig. S1B). This strong agreement was further supported by an intraclass correlation coefficient (ICC, type A-1 for absolute agreement, Arash Salarian 2008) of 0.982 (95% bootstrap CI: [0.980, 0.984]). These results indicate that both methods produce highly consistent centrosome size measurements across varying imaging conditions and experimental perturbations.

Comparison between 2D midplane measurements and full 3D segmentations showed systematic differences characterized by: 3D Radius = 1.053 · 2D Radius + 0.306 μm (95% CIs for slope: [1.046, 1.060]; intercept: [0.300, 0.312] μm; R^2^ = 0.977; Fig. S1C). This relationship indicates two key trends: a consistent positive intercept (0.306 μm), indicating that 2D measurements overrepresent the dense centrosomal core while underestimating the full 3D volume due to axial blurring; and a near-unity slope (1.053), demonstrating that proportional size changes are accurately captured regardless of absolute size. The high agreement (R^2^ = 0.977) supported the use of 2D measurements with a constant intercept correction for consistent size estimation.

We performed radial profile analysis on control and nocodazole-treated datasets using our established radial intensity profiling function. At the centrosome midplane and for the chromosome decondensation time frame, we generated symmetric 1D intensity distributions by computing the mean fluorescence intensity within concentrical elliptical rings aligned with the centrosome’s major and minor axes. The mean intensity profile and corresponding standard deviation for each experimental condition were calculated by averaging across anterior and posterior centrosomes from all embryos.

To quantify centrosome deformation, we calculated volumetric strain by comparing the spherical volume approximations of control and nocodazole-treated conditions. Centrosome volumes were estimated using V = 4/3πr^3^, where r represents the mean radius derived from intensity-based 2D measurements, corrected for intercepts. For each matched timepoint, strain was calculated as *ε* = (*Volume*_*Control*_ − *Volume*_*Nocodazole*_)/*Volume*_*Nocodazole*_. To account for measurement uncertainties, we propagated errors using first-order error analysis. The relative error in strain, *δε* was determined using 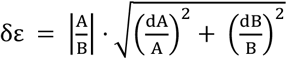 where A and B are the mean volumes of control and nocodazole-treated centrosomes and dA and dB are their respective standard deviations. To assess whether centrosome deformation scales with cell volume across developmental stages, we performed a weighted least-squares regression between mean cell volumes (derived from (122)) and centrosome strain (mean ± SD) measured in the time window −100 to −50 s relative to chromosome decondensation. Because the precision of strain measurements varied across cell stages, we assigned weights to each data point as the inverse of its squared measurement error (1/error^2^), ensuring that less precise estimates contributed less to the regression fit. The model took the form *ε* = *a* · *Cell volumes* + *b*, with parameters estimated via weighted linear regression (lscov in MATLAB). Confidence intervals (95%) for the slope (a) and intercept (b) were derived from the standard error of the estimates and a two-tailed t-distribution. Goodness-of-fit was evaluated using R^2^, and strict proportionality (strain ∝ volume) was tested by determining whether the 95% CI for the intercept included zero.

Centrosome aspect ratios were calculated from 2D midplane-segmented centrosomes using MATLAB’s regionprops function to extract the minor and major axis lengths, defined as: Aspect ratio = MinorAxisLength / MajorAxisLength. The orientation of centrosome deformation was quantified by calculating the angle between: 1) the inter-centrosome axis defined by a line connecting the 2D centroids of both centrosomes in the embryo midplane and 2) the major axis of an ellipse fitted to the segmented centrosome structure. For centrosome circularity calculations, measurements (from MATLAB’s regionprops) were corrected for pixelation artifacts at 63x magnification using an empirically-derived polynomial function. The correction accounts for how discrete pixel boundaries systematically reduce apparent circularity, especially for small centrosomes. The perfect circularity reference was calculated as: Perfect circle circularity = (0.9996 · r^2^ + 0.1771 · r + 0.4907) / r^2^ + -0.7655 · r + 0.185) where r is the equivalent radius. Coefficients were determined by measuring perfect circles across sizes. Final corrected values were obtained by dividing measured circularity by this reference.

Centrosome distance measurements were quantified in 3D space using the coordinates of both centrosomes and MATLAB’s pdist function. To assess the relative position of centrosomes to the embryo’s anteroposterior (AP) axis after metaphase onset (post-spindle assembly), we segmented the embryo shape using an intensity-based clustering approach (imsegkmeans in MATLAB using 4 clusters) and fitted an ellipse to the segmented embryo mask. The AP axis was defined as the major axis of this fitted ellipse, derived from its centroid, major/minor axis lengths, and orientation (MATLAB regionprops). For each centrosome, we computed its perpendicular distance to the AP axis by projecting its position onto the line defined by the ellipse’s major axis.

### Analysis of Spinning Disk Confocal Tubulin Image Sequences

Time synchronization was established using the onset of cortical contractility preceding cytokinetic ring ingression, with analysis restricted to post-metaphase events. GFP:TBB-2 images were corrected for camera background and photobleaching as for GFP:SPD-5.

For centrosome coordinate determination, we first segmented maximum intensity projections of GFP:TBB-2 z-stacks via intensity-based clustering (MATLAB imsegkmeans, 4 clusters) to approximate centrosome positions. The centrosome midplane was identified as the z-slice with maximal summed intensity within a 60-pixel diameter prismatic volume spanning all z-planes. We then refined coordinates by analyzing intensity profiles along 40 radial lines (MATLAB improfile), identifying boundary points where intensity gradients transitioned from positive to negative. Outlier removal was performed using Mahalanobis distance (MATLAB mahal). Final centrosome coordinates were derived from ellipse fits (MATLAB fitellipse) to boundary points (ellipse centroids), with quality control excluding ellipses showing aspect ratios <0.4 (minor/major axis) or mean squared error >180. From fitted ellipses, we generated 5-pixel (0.512 µm) thick binary masks (MATLAB poly2mask) and calculated the mean intensity across the mask thickness. Spindle axis alignment was achieved by: 1) defining the inter-centrosome line, 2) finding the two intersections with the ellipse, and 3) selecting the distal intersection point as the profile origin. Intensity profiles were then extracted clockwise from this origin within the masked regions, ensuring spatial alignment across embryos and time. The resulting intensity profiles were then subjected to linear interpolation (MATLAB’s interp1) to enforce consistent array size across all profiles.

Using the ellipse-derived centrosome coordinates, we generated concentric ring masks with a 30-pixel (3.07 µm) radius and a 20-pixel (2.05 µm) thickness to analyze microtubule organization in regions distal to the centrosome center. Intensity profiles were measured along radial lines spanning from the ring’s outer to inner perimeter (MATLAB improfile). These line profiles were averaged and the resulting mean intensity profile for each ring was aligned clockwise relative to the spindle axis as described above.

### Microtubule Volume Estimation

Using light microscopy and serial-section electron tomography data from Baumgart et al. (7), which measured a total polymerized tubulin concentration of ∼230 µM at centrosomes, we calculated microtubule occupancy within the PCM. For *C. elegans* microtubules (11 protofilaments, 20 nm diameter (44, 123)) containing 1354 tubulin dimers per micrometer, this concentration corresponds to 0.45 µm^3^ of microtubule volume. Within a metaphase centrosome volume of 14 µm^3^, this yields 3.21% PCM volume occupancy by microtubules.

### AFM Analysis

Force vs. indentation data recorded in QI mode and force volume maps were analyzed using a custom MATLAB pipeline (available at: https://github.com/Rufman91/ForceMapAnalysis). Prior to model fitting for indentation modulus estimations, force vs. indentation data were baseline corrected, applying both tilt (for drift correction) and offset adjustments, and the contact point determined using published methods (66, 114, 124). The Hertz-Sneddon model was fitted to the approach force-indentation data (excluding the lower 25% of data points) using a parabolic tip approximation derived from AFM tip and colloidal probe height data (125-127). Briefly, the indenter is described by a rotationally symmetric function *z* = *f*(*ρ*), which is rotated around the z-axis. The tip of the indenter is chosen to be *f*(*ρ* = 0) = 0 (with *f*(*ρ*) being indefinitely differentiable). When force F is applied, the indenter displaces into a half space by an amount of *δ*. For paraboloid indenters (as used for our nano-sized and colloidal probes), this produces a projected circle of contact at the surface with radius, *α*. In the region of contact, 0 < *ρ*/*α* < 1, the indenter shape is described by *z* = *f*(*x* = *ρ*/*α*) = *f*(*x*). Sneddon derived expressions for both the indentation *δ* (i.e., indenter’s displacement into the half space) and the applied force, *F* through simple integrals of the shape function.

The indentation, *δ* is:

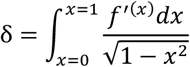

and the force, *F* is:

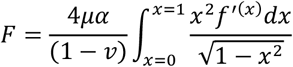

where *µ* is the shear modulus of the sample and *ν* is the Poisson’s ratio. For our nanosized tips and colloidal probes, this yields the following simple expression of force as a function of indentation depth:

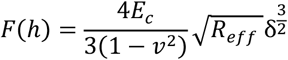

where *R*_*eff*_ is the effective radius of the sample and indenter in contact.

Here, *E*_*c*_ is the indentation modulus of the centrosome. The term *E*_*c*_/(1 − *ν*^2^) originates from the contact of two bodies with a considerable difference in their modulus of elasticity (centrosome and colloidal probe), where originally this term is the reduced modulus:

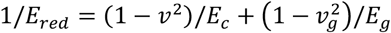

where *E*_*g*_ is the Young’s modulus of the colloid probe borosilicate glass (∼70 GPa) and *ν*_*g*_ is its Poisson’s ratio (0.2). Because *E*_*g*_ ≫ *E*_*c*_ the reduced modulus becomes: *E*_*red*_ = *E*_*c*_/(1 − *ν*^2^) To account for the centrosome’s radius of curvature, we implemented a geometrical correction. The effective radius *R*_*eff*_ between the tip and the sample was computed from smoothed contact height data.

Since the centrosome shape revealed areas with inclination towards the periphery and flatter areas towards the center, force curves for analysis were selected using to the following process. Contact height maps (height determined as the z-displacement at the contact point (128) underwent drift correction (using the surrounding glass height for baseline correction), smoothing and angle deconvolution to identify flat-top regions on each centrosome, applying an angular threshold of 1.45 radians (83.1°) (Fig. S4C). Force curve fits were quality-filtered using stringent criteria (Predictive R^2^ > 0.96; R^2^ > 0.99), with sub-threshold fits excluded from subsequent analysis (Fig. S4D-F). Centrosome size was determined from the contact height channel smoothed after manual segmentation of the centrosome volume.

The Hertz-Sneddon model assumes the sample to be a half space. However, the thickness of centrosomes is finite, varying between [126.7-979.6 nm]. To test putative effects of the finite thickness in experiments performed with the colloidal probe, thin film correction (not bonded, as the sample is attached but free to slide) was implemented (129). The corrected force vs. indentation expression for a not bonded thin film to the substrate is:

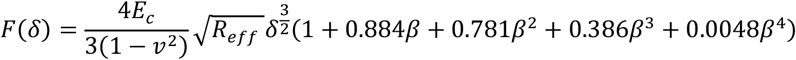

where *E*_*c*_ is the centrosome indentation modulus, *ν* is the centrosome’s Poisson’s ratio (taken as 0.5), and 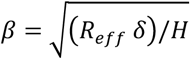, with *H* being the sample thickness (measured from contact height topography).

### Brillouin Image Quantification

Confocal GFP images were segmented using ilastik (v1.4.0b20post1) via pixel classification to identify centrosomes and chromosomes. A model was trained on multiple images representing different mitotic stages and subsequently applied in batch processing. The resulting segmentation files were converted to PNG masks in Fiji using a custom macro.

A custom MATLAB script processed these files, allowing for manual selection of centrosomes and chromosomes (labeled with the same fluorophore but clearly distinguishable by morphology and location) through user interaction, and generating separate centrosome and chromosome binary masks. The script rescaled Brillouin shift maps to match the GFP image dimensions using MATLAB’s ’imresize’ function (using bilinear interpolation) and applied overlapping pre-/post-image masks to the shift data, while enforcing a quality threshold (R^2^ > 0.8) for Lorentzian fits. For spindle space Brillouin shift quantification, we performed unsupervised cytoplasm segmentation using k-means clustering (8 clusters via imsegkmeans), selecting the cluster with the lowest mean intensity as the spindle space measurement. Mean values were calculated separately for (the two) centrosomes, chromosomes, and spindle space within each embryo.

To confirm that our centrosome masks captured true centrosome Brillouin shifts rather than boundary artifacts, we analyzed movement trajectories using MATLAB. Centrosome coordinates from pre- and post-scan confocal images (Fig. S5F) were used to model linear displacement during Brillouin scanning. We computed position probabilities during Brillouin acquisition, with earlier timepoints weighted toward the initial position and later ones toward the final position. The highest-probability position (designated position 0) defined the reference centrosome center. From this reference we generated line profiles along the movement trajectory, integrating Brillouin shifts across the centrosome diameter to derive mean values. All profiles were standardized with cytoplasm on the left and spindle space on the right (user-defined orientation). Trajectories were filtered to retain only those aligned with the spindle axis (±45° angle tolerance), excluding misaligned measurements.

### Nanolive Image Analysis

Fluorescence images were similarly segmented in ilastik (v1.4.0b20post1). A custom MATLAB script generated separate centrosome and chromosome binary masks, which were multiplied against the refractive index (RI) tomogram midplane. The spindle region was manually defined using a modified CROIEditor.m function (Jonas Reber, 2011) and similarly applied to the RI tomogram midplane (corresponding to the fluorescence imaging plane). Mean RI values were calculated for each compartment (centrosomes, chromosomes, and spindle space) per embryo.

### Longitudinal Modulus Calculation

The longitudinal modulus *M*^′^ was determined using the relation:

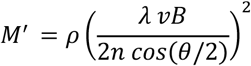

where *ρ* represents the mass density, *λ* is the laser wavelength, *θ* = 90° is the angle between the scattering and incident light, *νB* is the Brillouin shift and *n* is the refractive index (RI) from Nanolive holotomography. Sample density *ρ* was derived from RI measurements based on a two-component mixture model (68, 77) expressed as:

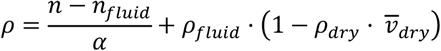

Here, *n*_*fluid*_ = 1.333 and *ρ*_*fluid*_ = 0.998 g/ml correspond to the refractive index and density of water, respectively, *α* = 0.190 ml/g represents the refraction increment for proteins and nucleic acids (68, 130-132) and 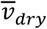 is the partial specific volume of the dry fraction. For most biological samples where 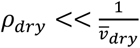, this simplifies to:

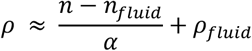

The longitudinal modulus *M*^′^ was calculated for each subcellular compartment (centrosomes, chromosomes, and spindle space) per embryo using a custom MATLAB script. This script used the mean Brillouin shift *νB* values per embryo with the corresponding compartment-specific mean mass density *ρ* and mean RI values derived from Nanolive holotomography analysis.

### Minimal Spring-Chain Model for Spindle Mechanics

To isolate and illustrate the physical effect of centrosome stiffness on spindle fluctuations, we constructed a minimal one-dimensional model of the spindle as a spring-mass chain (Fig. 6A). We represent the spindle as five mobile nodes, *x*_0_ through *x*_5_, bounded by fixed cortex positions at *x*_0_ = 0 (anterior) and *x*_6_ = *L* (posterior). The centrosomes are modeled as compliant springs (*k*_*c*_,*l*_*c*_) between nodes *x*_1_ − *x*_2_ (anterior) and *x*_4_ − *x*_5_ (posterior), connected to the cortex via fixed anterior (*k,l*_*ma*_) and posterior (*k,l*_*mp*_) cortical springs. The bridge region between the two centrosomes *x*_9_ − *x*_4_ consists of two identical springs (*k,l*_*s*_) centered on node *x*_3_, which approximates the chromosome location. Independent stochastic forces *F*_1_(*t*) and *F*_5_(*t*) act at the outer centrosomal edges, *x*_1_ and *x*_5_, modeling random fluctuations from the surrounding cytoplasm. Neglecting inertia, the dynamics follow overdamped Langevin equations:

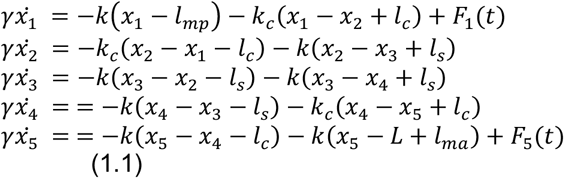

Here, *γ* is the viscous drag on each node, and *F*_1_(*t*), *F*_5_(*t*) are independent Gaussian white forces with a given variance *V*_0_. We adopt a value of *γ* = 150 *pN* · *s*/µ*m*, consistent with experimental estimates from centrosome centering dynamics in *C. elegans* embryos (133). We set *V*_0_ = 200 *pN*^2^*s*, but we note that our conclusions are robust to this choice, since altering *V*_0_ rescales the overall amplitude of fluctuations, but does not affect the observed trends. Bridge stiffness was set to *k* = 16 *pN*/µ*m*, consistent with effective values measured in *C. elegans* (133). For simplicity the same value was chosen for the cortical stiffness, whereas the centrosome stiffness *k*_*c*_ was scanned from 0.01*k* to 10*k*.

We simulated the overdamped dynamics of the six-spring chain (Eq. 1.1) using a forward Euler integration scheme with a timestep *d*_*t*_ = 0.01 *s*. Each simulation consisted of *N*_*s*_ = 100 stochastic trajectories of duration 110 s and simulated traces were downsampled to 10 Hz to mimic experimental frame rates (see below). For each centrosome stiffness *k*_*c*_, we calculated the temporal standard deviation of each observable, centrosome length σ(*x*_21_), spindle length σ(*x*_42_) and chromosome position σ(*x*_3_) for every trajectory and then averaged these values over the ensemble of simulations. All values used in simulations are summarized in Table S1.

### Power Spectral Density Analysis

To quantify how fluctuations are distributed in frequency we calculated one-sided power-spectral densities (PSDs) for the centrosome size *x*_21_(*t*) = *x*_2_(*t*) − *x*_1_(*t*) and the spindle length *x*_42_(*t*) = *x*_4_ (*t*) − *x*_2_(*t*).

For every trajectory, we first removed the temporal mean 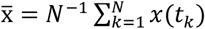 to eliminate any slow drift. With *N*_*f*_ = 550 points recorded at the experimental frame rate *f*_*s*_ = 10 Hz (Δt = 0.1 *s*) the discrete Fourier coefficients are:

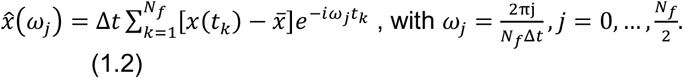

The one-sided PSD is then:

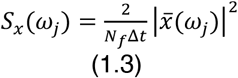

which we evaluated numerically with MATLAB’s Fast Fourier Transform (FFT). The factor of 2 folds negative frequencies onto positive ones, yielding units *µm*^2^/*Hz*. Spectra from 100 independent simulations were averaged to obtain an ensemble PSD for each *k*_*c*_/*k*.

Since absolute PSD heights depend on shot noise and calibration, we compared the PSD shapes, as they are more robust to the experimental unknowns. Thus, we used height-normalized PSDs, obtained by dividing each spectrum by its low frequency (*f* < 0.02 *Hz*), mean value. This normalization preserves knee positions and tail slopes while removing flat offsets.

### Mean-Squared Error

To determine which simulated PSD fits the experimental centrosome PSD best (Fig. 6D), we minimize the mean-squared error (MSE) over the frequency band (0.05 Hz-3 Hz), where biological signal dominates. Using logarithmic values:

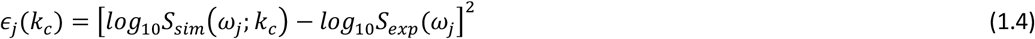

where *S*_*exp*_ (*ω*_*j*_) is the experimental power spectrum and *S*_*sim*_ (*ω*_*j*_ ; *k*_*c*_) is the simulated one for centrosome stiffness *k*_*c*_, we calculate the cost function for each stiffness:

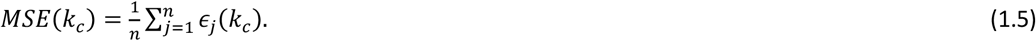

The *k*_*c*_ that yields the smallest MSE is our best estimate for the PCM stiffness.

### Power Spectral Density Analysis of Experimental *C. elegans* Movies

For image preprocessing, time-lapse movies of *C. elegans* embryos expressing GFP:SPD-5 (acquired at 10 Hz) were subjected to mild Gaussian filtering (kernel size: 1, σ = 1) to reduce noise. Binary segmentation was performed using Otsu’s adaptive thresholding (MATLAB’s multithresh), followed by hole-filling (imfill) to generate contiguous masks.

For centrosome tracking, an initial region of interest (ROI) was manually defined on the posterior centrosome using the maximum intensity projection over time with a modified version of the CROIEditor.m tool (Jonas Reber, 2011). For each frame, binary objects within the ROI were detected using MATLAB’s regionprops, sorted by area, and filtered to retain the largest valid structure. The posterior boundary coordinates were computed using MATLAB’s bwboundaries function. Two points on this boundary, an inner (closer to the anterior centrosome) and an outer point (farther from it), were identified by projecting the anterior-posterior (AP) axis onto the boundary. These points were validated by enforcing a minimum separation of half the object’s equivalent diameter.

For oscillatory analysis, we focused on a 110 second time window (−90 to +20 s relative to cytokinesis onset), where spindle oscillations are most prominent (Fig. 6C). The posterior centrosome length (distance between inner and outer boundary points) was computed for each frame. To analyze these dynamics in the frequency domain, we calculated the one-sided power spectral density (PSD) using a custom MATLAB function. The function first applied a fast Fourier transform (FFT MATLAB’s function) to the distance signal while removing the DC component (mean value) to isolate fluctuations. The resulting two-sided power spectrum was then converted to a one-sided representation by retaining only the positive frequencies and doubling positive frequencies except DC (zero). The frequency axis was scaled based on the 10 Hz sampling rate and the PSD values normalized to a 1-Hz frequency bandwidth for comparison.

### Statistical Analysis

Statistical analysis was performed using a custom workflow in MATLAB (R2021b). Each dataset was first tested for normal distribution using the Lilliefors test. The homogeneity of variances was assessed using Levene’s test. For comparisons involving two groups, if both datasets were normally distributed, an unpaired, two-tailed Student’s t-test was used. If the data were normal but variances were unequal, Welch’s t-test was applied. If the data failed the normality test for at least one of the groups, a Mann-Whitney U test was used. For comparisons of more than two groups, a one-way ANOVA was used if all datasets were normal and had equal variances. If the data were non-normal or had unequal variances, a Kruskal-Wallis test was employed. If a significant main effect was found (p < 0.05) for comparisons of more than two groups, post-hoc analyses were conducted: the Tukey-Kramer test was used following ANOVA or Bonferroni-corrected pairwise comparisons were performed following the Kruskal-Wallis test. For all experiments, the significance threshold was taken as p < 0.05. Significance levels are defined as follows: ****p < 0.0001, ***p < 0.001, **p < 0.01, *p < 0.05; n.s., not significant.

Analysis of centrosome volume and fluorescence intensity over time was performed using a linear mixed-effects model in MATLAB (fitlme) to account for the non-independence of repeated measurements collected from the same embryo across multiple time points. The model was specified as Volume (or Intensity) ∼ Group · Time + (1 | Embryo). This structure assessed the fixed effects of experimental group, time, and their interaction, while the random intercept (1 | Embryo) accounted for the inherent variability between individual embryos. Significance of the fixed effects was assessed at α = 0.05.

Distance correlation analysis (distcorr, MATLAB; Shen Liu, 2013), a nonparametric test, was used to assess the potential nonlinear relationship between centrosome stiffness (indentation modulus) and size. Statistical significance (α = 0.05) was determined through permutation testing (10,000 iterations), which created a null distribution by randomly shuffling stiffness values against size measurements. The resulting p-value represents the proportion of these random permutations that showed a correlation strength equal to or greater than the observed relationship, confirming that the association was unlikely to occur by chance.

## Acknowledgments

We would like thank members of the Dammermann and Campbell laboratories, Oliver Paulin and David Zwicker (MPI-DS Göttingen) for discussions; the *Caenorhabditis* Genetics Center (CGC), Antony Jose (University of Maryland) and Nathalie Pujol (CIML Marseille) for strains; Carsten Janke (Institut Curie, Paris), Julien Dumont and Benjamin Lacroix (Institut Jacques Monod, Paris) for help with preparation of purified tubulin; Tamara Tomin and Ruth Birner-Gruenberger (TU Wien) for allowing us to use their Nanolive microscope; and Josef Gotzmann, Thomas Peterbauer and Clara Bodner of the Max Perutz Labs BioOptics Facility for technical assistance. This work was supported by funding from the Austrian Science Fund FWF, grants P34526-B and F8803-B to A. Dammermann, as well as a Max Perutz PhD fellowship of the Max Perutz Labs to J. Garcia-Baucells. C. Rumpf-Kienzl was the recipient of an INDICAR post-doctoral fellowship, funded by the Mahlke-Obermann Stiftung and the European Union Seventh Framework Programme under grant agreement 609431. O.G. Andriotis, P.J. Thurner and M. Rufin acknowledge funding by the Vienna Science and Technology Fund (WWTF), grant 10.47379/LS19035. S. Fürthauer and A. Zampetaki were supported by the Vienna Science and Technology Fund (WWTF), grant 10.47379/VRG20002. Calculations were performed using supercomputer resources provided by the Austrian Scientific Computing Infrastructure (ASC). R. Prevedel acknowledges support of an ERC Consolidator Grant (no. 864027, Brillouin4Life) and the German Center for Lung Research (DZL). This work was supported by funds from the European Molecular Biology Laboratory.

## Notes

### Competing Interest Statement

The authors have declared no competing interest.

