## Supplemental Material for "Centrosome Softening As A Mechanical Adaptation For Mitosis"

**INDEX OF SUPPLEMENTAL MATERIAL**

**SUPPLEMENTAL FIGURES:**

Figure S1: Reference measurements of centrosome morphology in control and nocodazole-treated embryos.
Figure S2: Further characterization of the impact of cortical forces on centrosome deformation.
Figure S3: Further characterization of KLP-7 depletion.
Figure S4: AFM analysis of isolated centrosomes.
Figure S5: Further characterization of one-cell embryos by Brillouin microscopy and holotomography.
Figure S6: Further characterization of *C. elegans* embryos by Brillouin microscopy and holotomography.

**SUPPLEMENTAL TABLES:**

Table S1: Key model parameters and their provenance.
Table S2: Timing of events in the one-cell *C. elegans* embryo aligned to three different reference time points.

### Figure S1

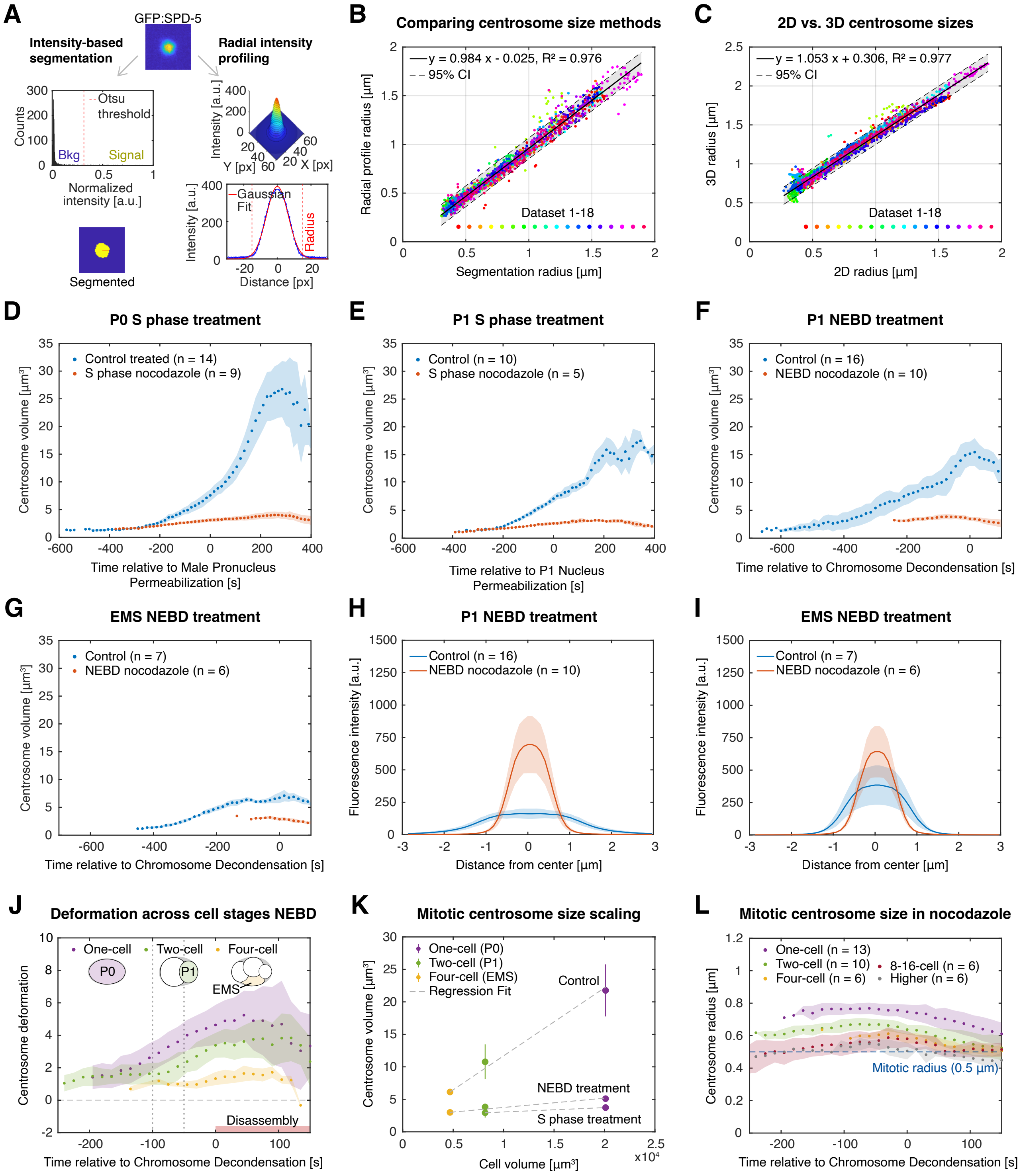
A
GFP:SPD-5
Intensity-based segmentation
Radial intensity profiling
Otsu threshold
Counts
Bkg
Signal
Normalized intensity [a.u.]
Segmented
Intensity [a.u.]
Y [px]
X [px]
Gaussian Fit
Radius
Distance [px]
B
Comparing centrosome size methods
y = 0.984 x - 0.025, R² = 0.976
95% CI
Dataset 1-18
Radial profile radius [µm]
Segmentation radius [µm]
C
2D vs. 3D centrosome sizes
y = 1.053 x + 0.306, R² = 0.977
95% CI
Dataset 1-18
3D radius [µm]
2D radius [µm]
D
P0 S phase treatment
Control treated (n = 14)
S phase nocodazole (n = 9)
Centrosome volume [µm³]
Time relative to Male Pronucleus Permeabilization [s]
E
P1 S phase treatment
Control (n = 10)
S phase nocodazole (n = 5)
Time relative to P1 Nucleus Permeabilization [s]
F
P1 NEBD treatment
Control (n = 16)
NEBD nocodazole (n = 10)
Time relative to Chromosome Decondensation [s]
G
EMS NEBD treatment
Control (n = 7)
NEBD nocodazole (n = 6)
Time relative to Chromosome Decondensation [s]
H
P1 NEBD treatment
Control (n = 16)
NEBD nocodazole (n = 10)
Fluorescence intensity [a.u.]
Distance from center [µm]
I
EMS NEBD treatment
Control (n = 7)
NEBD nocodazole (n = 6)
Distance from center [µm]
J
Deformation across cell stages NEBD
One-cell
Two-cell
Four-cell
P0
P1
EMS
Disassembly
Centrosome deformation
Time relative to Chromosome Decondensation [s]
K
Mitotic centrosome size scaling
One-cell (P0)
Two-cell (P1)
Four-cell (EMS)
Regression Fit
Control
NEBD treatment
S phase treatment
Centrosome volume [µm³]
Cell volume [µm³]
x10⁴
L
Mitotic centrosome size in nocodazole
One-cell (n = 13)
Two-cell (n = 10)
Four-cell (n = 6)
8-16-cell (n = 6)
Higher (n = 6)
Mitotic radius (0.5 µm)
Centrosome radius [µm]
Time relative to Chromosome Decondensation [s]

**Figure S1. Reference measurements of centrosome morphology in control and nocodazole-treated embryos (related to Fig. 1).** (A) Alternative methods for centrosome size measurement from 2D midplane images (GFP:SPD-5). Left: Intensity-based segmentation using Otsu thresholding, which automatically separates signal (above threshold) from background (below). The resulting binary mask is used to calculate an equivalent radius (red line) from a circle of equal area. Right: Radial intensity profiling uses concentric elliptical rings (aligned with the centrosome's centroid and orientation) to extract mean intensity values within each ring. The mirrored radial profile is fitted with a Gaussian function and the full-width half-maximum (FWHM) is converted to a physical radius. (B) Comparison of centrosome size estimations from 2D midplane images across multiple datasets representing different biological perturbations and imaging conditions by intensity-based segmentation and radial intensity profiling shows close agreement between the two methods. (C) Comparison of centrosome size estimations by intensity-based segmentation in 2D and 3D datasets reveals close agreement with a systematic 0.3 µm offset (2D radii smaller). This offset arises because the 2D midplane overrepresents the dense core while missing the faint, axially blurred periphery captured in 3D. The offset is size-independent and primarily reflects the microscope's fixed point spread function (PSF). Data represent 18 datasets, comprising 5-25 embryos per dataset. Datapoints are centrosome means averaged across embryos per dataset. (D-G) Nocodazole treatment at S phase or nuclear envelope breakdown (NEBD) strongly impacts mitotic centrosome expansion in one- (P0), two- (P1) and four-cell stage (EMS) embryos, albeit to a lesser extent in later cell stages. (H, I) 2D radial profiling of GFP:SPD-5 fluorescence intensities in nocodazole-treated embryos in mitosis shows the PCM scaffold increases in density close to the centrioles in two- (P1) and four-cell stage (EMS) embryos. Control and nocodazole radial profiles become more similar at later cell stages. (J) PCM deformation assessed during mitosis in one-, two- and four-cell stage embryos shows less deformation in subsequent divisions. Nocodazole treatment at NEBD was used as a baseline (one-cell data from Fig. 1B; two-cell from Fig. S1F; four-cell from Fig. S1G). (K) Mean centrosome volume (measured during the mitotic time window indicated by dotted lines in J) scales with cell volume only in untreated embryos. Nocodazole treatment in S phase or mitosis essentially eliminates size scaling. (L) Mitotic centrosome size in the absence of microtubules (i.e., upon nocodazole treatment at NEBD) consistently exceeds 0.5 µm in radius across early developmental stages (important for cell cycle stage assignment of isolated centrosomes in AFM experiments; Fig. 4). Points/solid lines in D-L indicate the mean of the two centrosomes in each embryo across embryos; error bars or shading represent the associated standard deviation.

### Figure S2

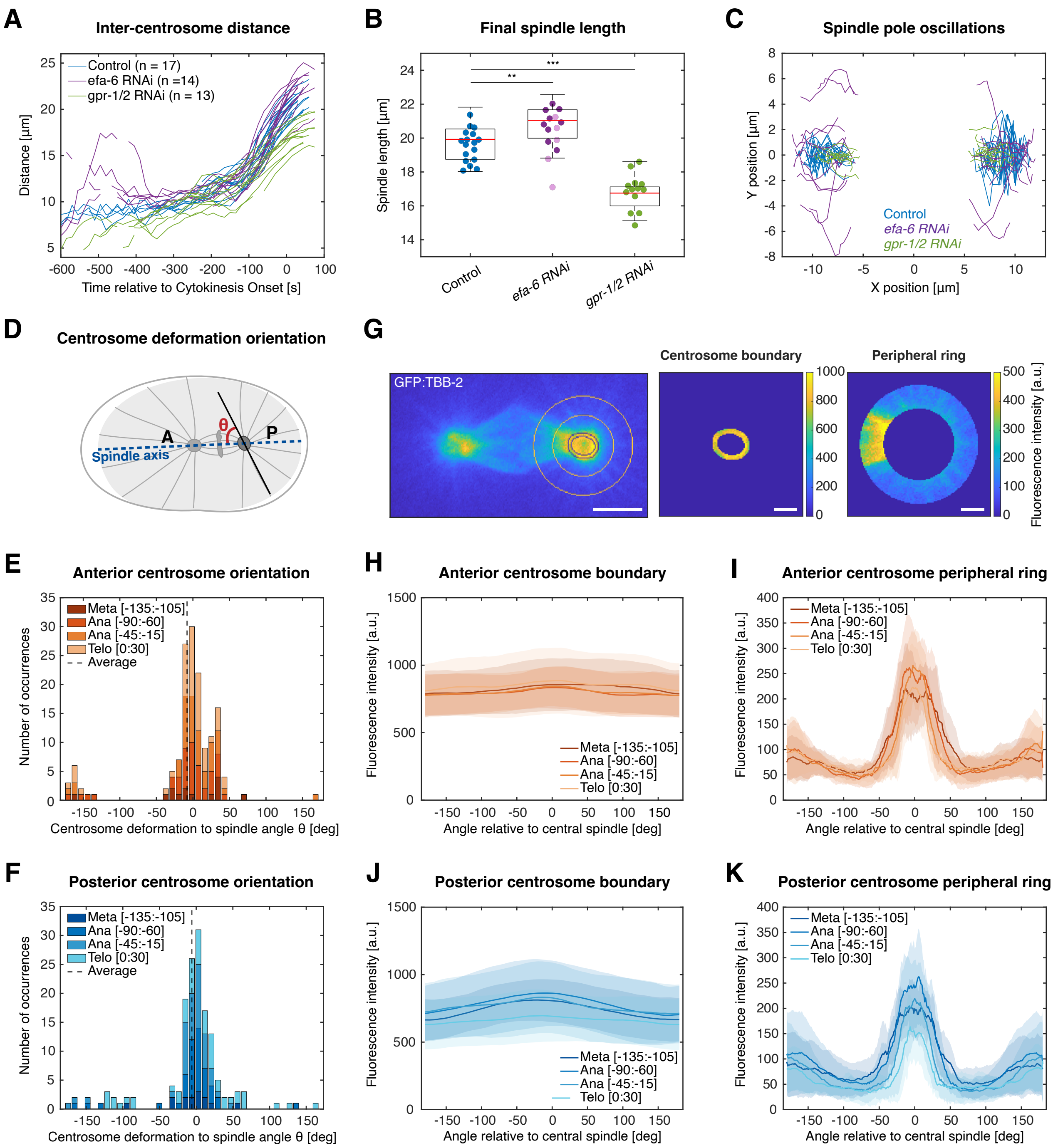
A
Inter-centrosome distance
Control (n = 17)
efa-6 RNAi (n =14)
gpr-1/2 RNAi (n = 13)
Distance [µm]
Time relative to Cytokinesis Onset [s]
B
Final spindle length
Spindle length [µm]
Control
efa-6 RNAi
gpr-1/2 RNAi
C
Spindle pole oscillations
Y position [µm]
X position [µm]
Control
efa-6 RNAi
gpr-1/2 RNAi
D
Centrosome deformation orientation
A
P
θ
Spindle axis
G
GFP:TBB-2
Centrosome boundary
Peripheral ring
Fluorescence intensity [a.u.]
E
Anterior centrosome orientation
Meta [-135:-105]
Ana [-90:-60]
Ana [-45:-15]
Telo [0:30]
Average
Number of occurrences
Centrosome deformation to spindle angle θ [deg]
H
Anterior centrosome boundary
Fluorescence intensity [a.u.]
Angle relative to central spindle [deg]
I
Anterior centrosome peripheral ring
F
Posterior centrosome orientation
J
Posterior centrosome boundary
K
Posterior centrosome peripheral ring

**Figure S2. Further characterization of the impact of cortical forces on centrosome deformation (related to Fig. 2).** (A-C) Inter-centrosome distance (A), spindle length at cytokinesis onset (B) and anaphase spindle pole oscillations (C) in controls, *efa-6(RNAi)* and *gpr-1/2(RNAi)* embryos. *efa-6(RNAi)* embryos display centrosome detachment from the male pronucleus resulting in increased inter-centrosome distances during pronuclear migration, increased spindle lengths ($\Delta$ = 1.51 μm, p = 0.007; light purple datapoints indicating embryos with transversely aligned spindles excluded from one-way ANOVA/Tukey-Kramer analysis), impaired anaphase spindle oscillations and spindle alignment defects (52). *gpr-1/2(RNAi)* embryos display reduced spindle length ($\Delta$ = -3.13 μm, p = 1.97e-08) and an absence of anaphase spindle oscillations, as previously reported (52-54). (D-F) Centrosome deformation primarily aligns with the spindle axis during mitosis. Schematic (D) and measurements of the main axis of deformation for anterior (E) and posterior (F) centrosomes visualized by GFP:SPD-5, defined as the angle between the long axis of fitted centrosome ellipses and the spindle axis at indicated mitotic stages (numbers in brackets indicate time window per stage in s). (G-K) Microtubules emanate in all directions from the centrosome boundary but are enriched towards the central spindle further away from the centrosome. Schematic (G) and measurement of tubulin fluorescence levels (GFP:TBB-2) at anterior and posterior centrosomes at the centrosome boundary (H, J) and 3-5 μm from the centrosome center (I, K). Scale bars: 5 μm (overview), 2 μm (enlarged images), respectively. Error bars or shading in B, E, F, H-K represent standard deviation. Sample sizes: (E, F) n = 17 embryos; (H- K) n = 16 embryos.

### Figure S3

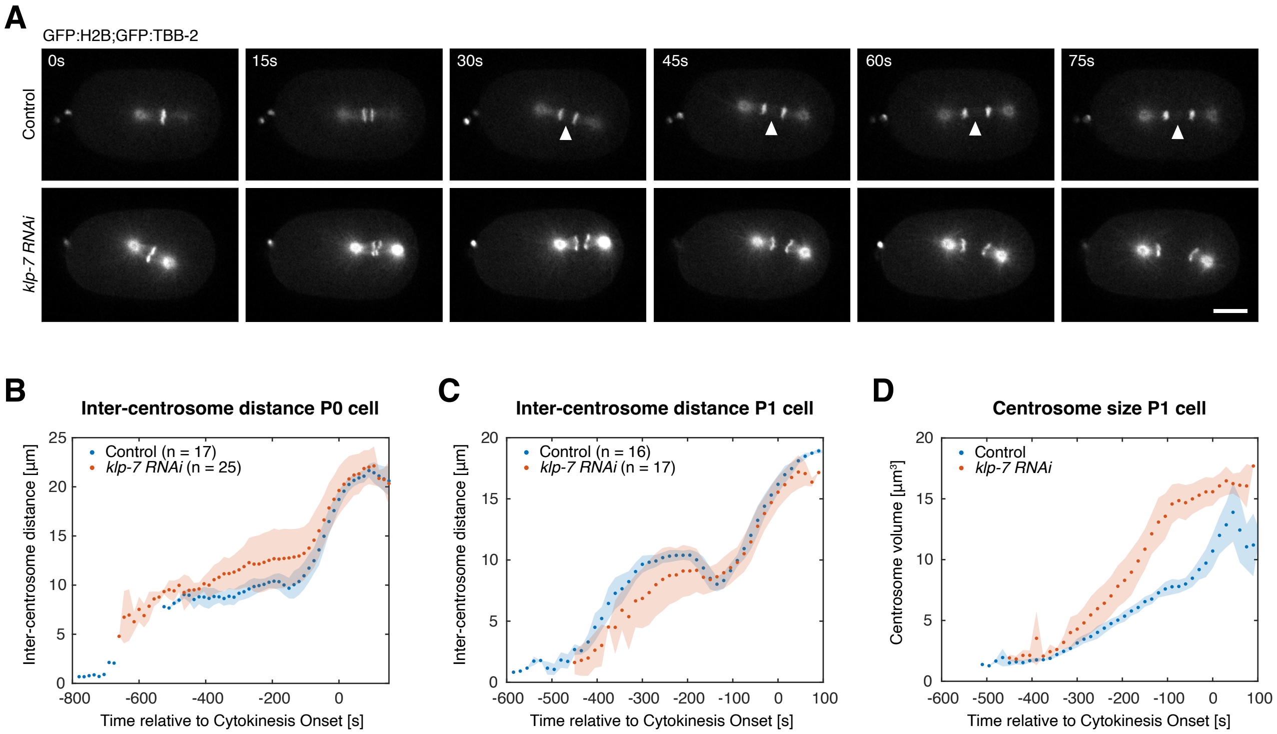
A
GFP:H2B;GFP:TBB-2
0s
15s
30s
45s
60s
75s
Control
klp-7 RNAi
B
Inter-centrosome distance P0 cell
Control (n = 17)
klp-7 RNAi (n = 25)
Inter-centrosome distance [µm]
Time relative to Cytokinesis Onset [s]
C
Inter-centrosome distance P1 cell
Control (n = 16)
klp-7 RNAi (n = 17)
Inter-centrosome distance [µm]
Time relative to Cytokinesis Onset [s]
D
Centrosome size P1 cell
Control
klp-7 RNAi
Centrosome volume [µm³]
Time relative to Cytokinesis Onset [s]

**Figure S3. Further characterization of KLP-7 depletion (related to Fig. 3).** (A) KLP-7 depletion in embryos co-expressing GFP:histone H2B and GFP: β-tubulin results in increased centrosomal microtubule levels compared to controls and spindle breakage during anaphase, as previously reported (58, 59). Note also the loss of central spindle microtubules found in controls (arrowheads). Scale bar: 10 μm. (B) KLP-7 depletion in one-cell stage embryos results in increased inter-centrosome distances prior to anaphase onset, as assessed in embryos expressing GFP:SPD-5. (C) Inter-centrosome distances are decreased in the two-cell stage P1 cell, as previously reported (60). (D) Centrosome size is also increased in two-cell stage (P1) embryos. In B-D, points represent the mean of anterior/posterior centrosomes (averaged per embryo) across embryos; shading shows standard deviation.

### Figure S4

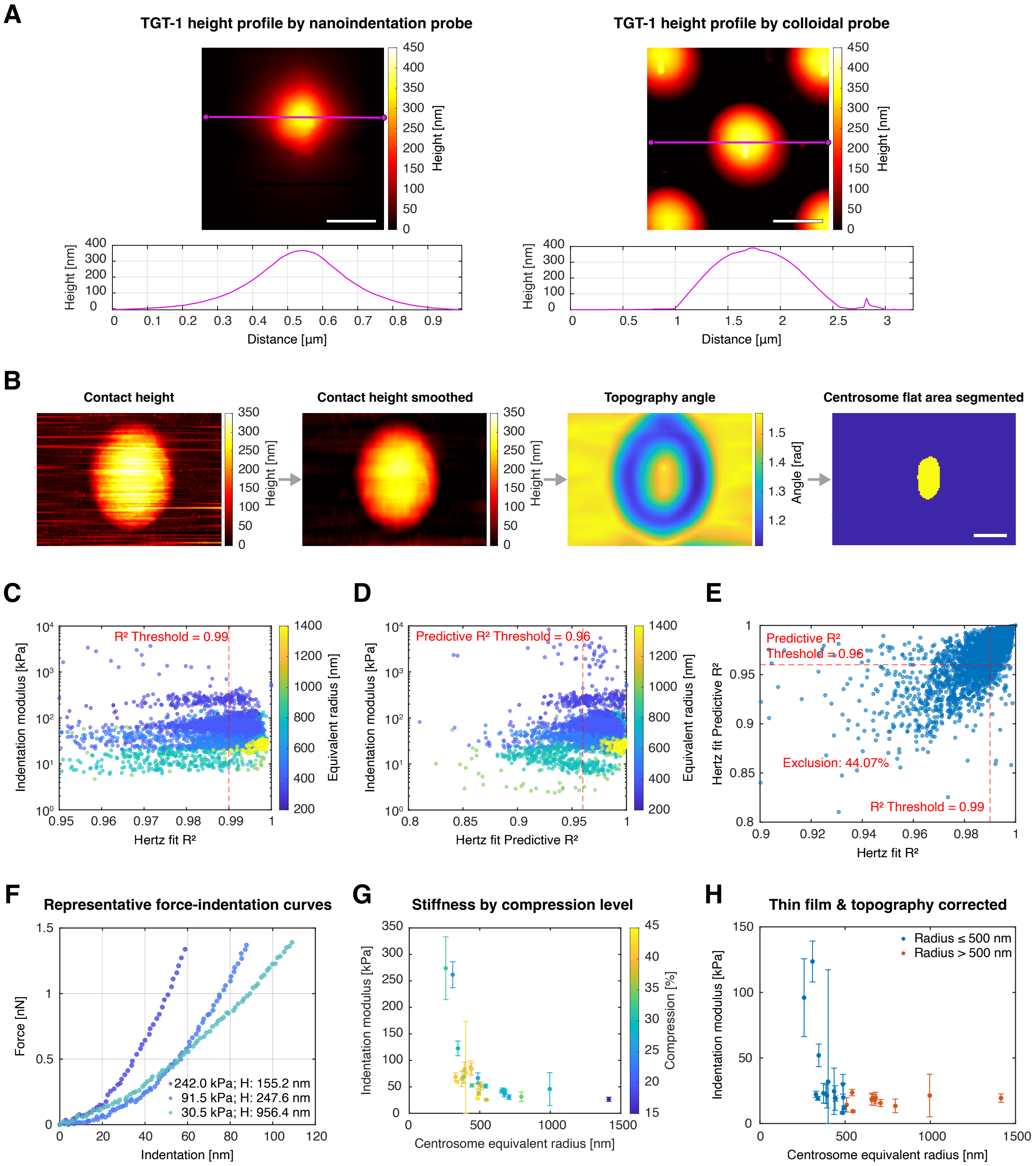

A
TGT-1 height profile by nanoindentation probe
TGT-1 height profile by colloidal probe
Height [nm]
Distance [µm]
B
Contact height
Contact height smoothed
Topography angle
Centrosome flat area segmented
Angle [rad]
C
R² Threshold = 0.99
Indentation modulus [kPa]
Equivalent radius [nm]
Hertz fit R²
D
Predictive R² Threshold = 0.96
Hertz fit Predictive R²
E
Predictive R² Threshold = 0.96
Exclusion: 44.07%
R² Threshold = 0.99
F
Representative force-indentation curves
Force [nN]
Indentation [nm]
242.0 kPa; H: 155.2 nm
91.5 kPa; H: 247.6 nm
30.5 kPa; H: 956.4 nm
G
Stiffness by compression level
Compression [%]
Centrosome equivalent radius [nm]
H
Thin film & topography corrected
Radius ≤ 500 nm
Radius > 500 nm

**Figure S4. AFM analysis of isolated centrosomes (related to Fig. 4).** (A) Prior to mechanical testing, AFM tip geometry was characterized through reverse reconstruction using TGT1 calibration gratings for both nanoindentation and colloidal bead probes (114). Scale bar: 1 μm. (B) Workflow to identify flat centrosome regions for micromechanical analysis using colloidal probe. Scale bar: 0.5 μm. (C-E) Quality-filtering of force curve fits, showing thresholds for exclusion from subsequent analysis (Predictive $R^2 > 0.96$; $R^2 > 0.99$). Note that filtering does not introduce bias. (F) Representative examples of force indentation curves across the stiffness spectrum. (G) Degree of compression does not correlate with stiffness. (H) Thin film (not bonded) and curvature corrections do not eliminate the size-stiffness relationship. Error bars in G, H represent standard deviation. Sample size: n = 27 centrosomes.

### Figure S5

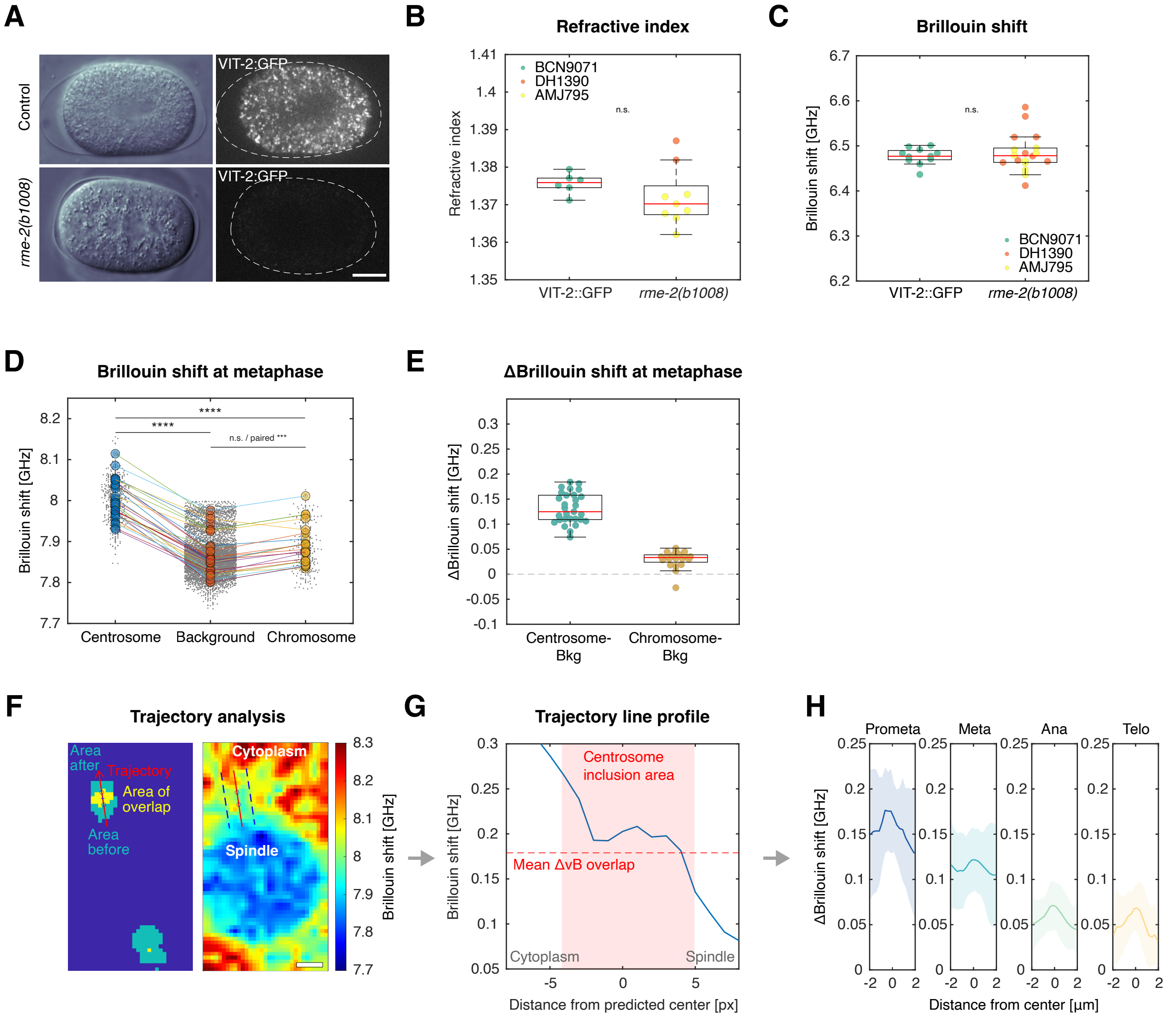
A
Control
rme-2(b1008)
VIT-2:GFP
VIT-2:GFP
B
Refractive index
BCN9071
DH1390
AMJ795
n.s.
Refractive index
VIT-2::GFP
rme-2(b1008)
C
Brillouin shift
n.s.
Brillouin shift [GHz]
BCN9071
DH1390
AMJ795
VIT-2::GFP
rme-2(b1008)
D
Brillouin shift at metaphase
****
****
n.s. / paired ***
Brillouin shift [GHz]
Centrosome
Background
Chromosome
E
ΔBrillouin shift at metaphase
ΔBrillouin shift [GHz]
Centrosome-Bkg
Chromosome-Bkg
F
Trajectory analysis
Area after
Trajectory
Area of overlap
Area before
Cytoplasm
Spindle
Brillouin shift [GHz]
G
Trajectory line profile
Centrosome inclusion area
Mean ΔvB overlap
Cytoplasm
Spindle
Brillouin shift [GHz]
Distance from predicted center [px]
H
Prometa
Meta
Ana
Telo
ΔBrillouin shift [GHz]
Distance from center [μm]

**Figure S5. Further characterization of one-cell embryos by Brillouin microscopy and holotomography (related to Fig. 5).** (A) *rme-2* mutant embryos are deficient in yolk granules, here marked by VIT-2:GFP. Scale bar: 10 µm. (B, C) Refractive index measurements by holotomography (B; control n = 6, *rme-2(b1008)* n = 9; Welch's t-test, p = 0.2316) (B) and Brillouin shift (C; control n = 11, *rme-2(b1008)* n = 18; Welch's t-test, p = 0.5144) (C) reveal no differences in cytoplasmic signal between *rme-2* yolk-deficient mutants and wild-type controls. (D, E) Derivation of relative Brillouin frequency shift of centrosomes and chromosomes for a representative dataset. Plot in (D) shows raw pixel Brillouin shift values (dots) and averages for each area (filled circles) at metaphase, with lines connecting measurements within a single embryo (centrosome n = 28, spindle background n = 28, chromosome n = 20 embryos; Kruskal-Wallis, $p < 0.0001$; post-hoc Bonferroni: centrosome vs. background p = 2.964e−10; centrosome vs. chromosome p = 1.033e−05; background vs. chromosome p = 0.620, although chromosome values exceed paired background (p = 0.00019, Wilcoxon). Plot in (E) shows relative frequency shift normalized to each embryo's spindle background for the same dataset at metaphase. (F-H) Trajectory analysis rules out boundary effects for centrosome measurements. (F) Example Brillouin frequency shift map and image masks showing centrosome position at prometaphase before/after Brillouin scan, defining its trajectory during acquisition. Scale bar: 2 µm. (G) Line scan over the centrosome trajectory showing the normalized Brillouin frequency shift over background for the example image shown in F. (H) Average relative Brillouin frequency line profiles for centrosomes generated as in F, G at indicated mitotic stages (prometaphase: n = 19; metaphase: n = 29; anaphase: n = 22; telophase: n = 14). In B-E, boxplots show the median (red line), IQR (box), and range. Colored data points represent the mean per region of interest. In H, shaded area represents the standard deviation and the solid line represents the mean.

### Figure S6

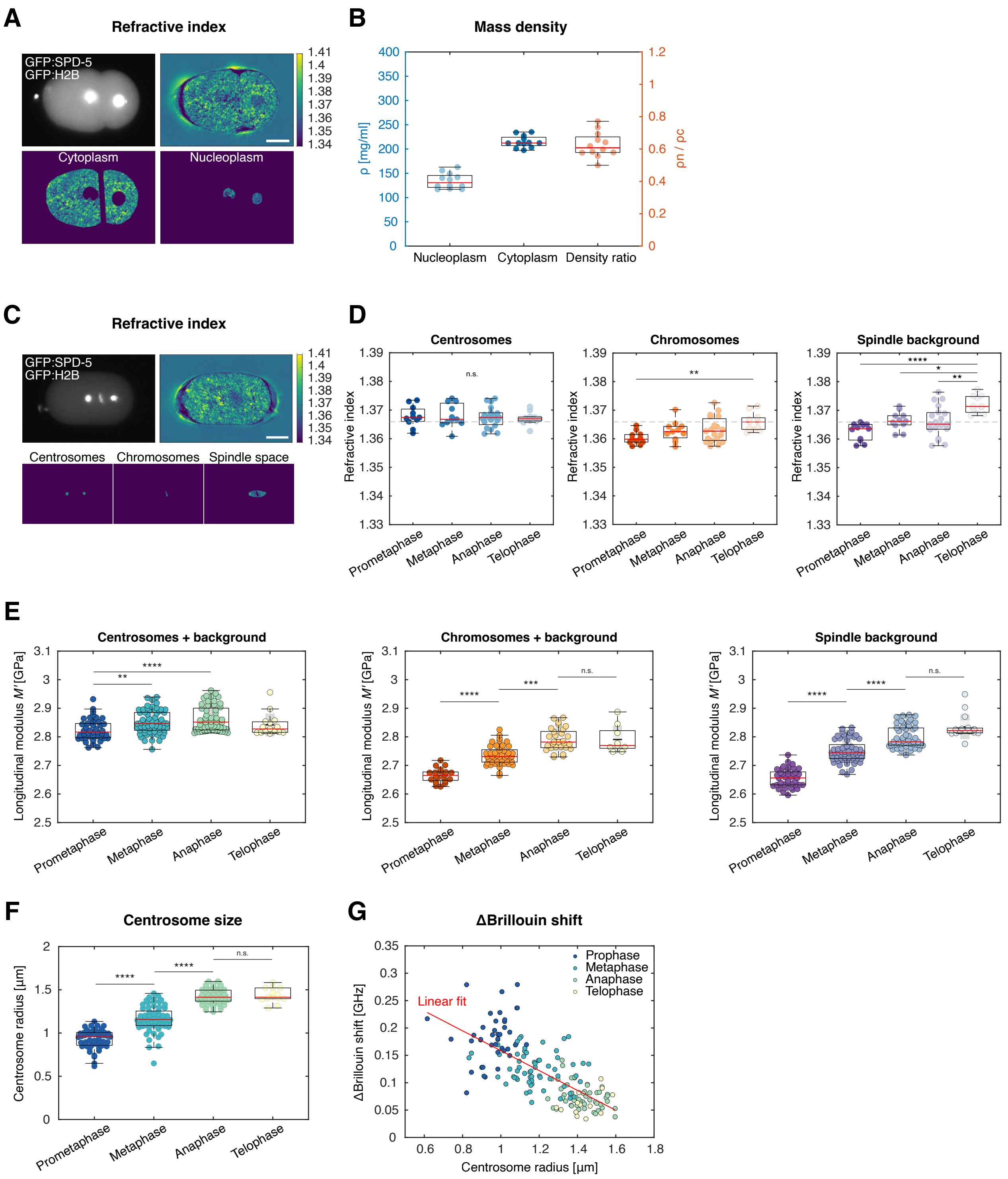

A
Refractive index
GFP:SPD-5
GFP:H2B
Cytoplasm
Nucleoplasm
B
Mass density
ρ [mg/ml]
ρn / ρc
Nucleoplasm
Cytoplasm
Density ratio
C
Refractive index
Centrosomes
Chromosomes
Spindle space
D
Centrosomes
Chromosomes
Spindle background
Refractive index
Prometaphase
Metaphase
Anaphase
Telophase
E
Centrosomes + background
Chromosomes + background
Spindle background
Longitudinal modulus M′ [GPa]
F
Centrosome size
Centrosome radius [µm]
G
ΔBrillouin shift
ΔBrillouin shift [GHz]
Centrosome radius [µm]
Prophase
Metaphase
Anaphase
Telophase
Linear fit

**Figure S6. Further characterization of *C. elegans* embryos by Brillouin microscopy and holotomography (related to Fig. 5).** (A, B) Correlative holotomography and fluorescence imaging of interphase embryos at one-, two- and four-cell stages reproduces established mass density relationships between nucleus and cytoplasm (77), with nucleoplasmic density lower than that of the cytoplasm. (A) Example interphase two-cell stage embryo showing fluorescence and refractive index measurements. Scale bar: 10 µm. (B) Computed mass density of each compartment and nucleocytoplasmic ratio for individual embryos (n = 13 embryos, 1-4 cell stage, DAM1342 strain). (C, D) Refractive index measurements for centrosomes, chromosomes and spindle space at indicated mitotic stages show significant increase in refractive index for chromosomes and spindle space, but no change for centrosomes. (C) Example of fluorescence and refractive index measurement of a metaphase one-cell stage embryo with masks defining centrosomes, chromosomes and spindle space. Scale bar: 10 µm. (D) Refractive index measurements across mitotic stages based on images as in B (dashed line: RI = 1.3659, cytoplasmic refractive index; (77, 119). Centrosomes remain stable (prometaphase: n = 11; metaphase: n = 11; anaphase: n = 19; telophase: n = 11; one-way ANOVA, p = 0.910). Chromosomes show stage-dependent changes (same n; ANOVA, p = 0.0075; post-hoc Tukey-Kramer: telophase vs. prometaphase, p = 0.0035; other comparisons p > 0.16). Spindle background increases progressively (same n; ANOVA, p < 0.0001; post-hoc: prometaphase-telophase p = 1.0296e−05; metaphase-telophase p = 0.012; anaphase-telophase p = 0.0024; early stages p > 0.10). (E) Longitudinal modulus calculation shows an increase in elasticity for chromosomes and spindle space from prometaphase to telophase, potentially due to spindle envelope disassembly, while centrosomes maintain near-constant elasticity across mitosis despite a stiffening environment. Centrosomes show significant variation (prometaphase n = 42, metaphase n = 63, anaphase n = 53, telophase n = 15; Kruskal-Wallis, p = 0.0001; post-hoc Bonferroni: prometaphase-metaphase p = 0.0029; prometaphase-anaphase p = 9.614e−05; later stages p > 0.54). Chromosomes exhibit strong stage-dependence (prometaphase n = 20, metaphase n = 45, anaphase n = 29, telophase n = 9; Kruskal-Wallis, p < 0.0001; post-hoc: prometaphase-metaphase p = 5.364e−05; metaphase-anaphase p = 0.00014; anaphase-telophase p = 1.0). Spindle background stiffness increases progressively (prometaphase n = 40, metaphase n = 61, anaphase n = 53, telophase n = 15; Kruskal-Wallis, p < 0.0001; post-hoc: prometaphase-metaphase p = 3.009e−08; metaphase-anaphase p = 5.497e−05; anaphase-telophase p = 0.592). (F, G) Relative Brillouin shift of centrosomes negatively correlates with size as mitosis progresses in one-cell embryos. (F) Centrosome radius derived from fluorescence confocal measurements on the Brillouin setup (prometaphase: n = 50; metaphase: n = 67; anaphase: n = 53; telophase: n = 14, Kruskal-Wallis, p < 0.0001; post-hoc Bonferroni: prometaphase-metaphase p = 8.554e−07, metaphase-anaphase p = 1.180e−09, anaphase-telophase p = 1.0). (G) Correlation of relative Brillouin shift with centrosome size (n = 37 centrosomes; linear fit: y = -0.18x + 0.34, r = -0.72, p = 9.5e-29). In B, D, E and F, boxplots show the median (red line), IQR (box), and range. Colored datapoints represent the mean per region of interest.

**Table S1. Key model parameters and their provenance.**

| Parameter | Symbol | Value | Units | Source |
|---|---|---|---|---|
| Effective cytoplasmic drag | $\gamma$ | 150 | pN s $\mu m^{-1}$ | (133) |
| Effective spindle spring constant | $k$ | 16 | pN $\mu m^{-1}$ | (133) |
| PCM spring constant (best fit) | $k_c$ | 1.53 | pN $\mu m^{-1}$ | Fitted (log-MSE fit) |
| Noise power | $V_0$ | 200 | $pN^2$ s | Assumed/fixed for convenience |
| Average cell length | $L$ | 51.4 | μm | Measured (this study) |
| Average centrosome length | $l_c$ | 2.7 | μm | Measured (this study) |
| Average centrosome-chromosome distance | $l_s$ | 8 | μm | Measured (this study) |
| Posterior cortex-centrosome distance | $l_{mp}$ | 10 | μm | Measured (this study) |
| Anterior cortex-centrosome distance | $l_{ma}$ | 20 | μm | Measured (this study) |

**Table S2. Timing of events in the one-cell *C. elegans* embryo aligned to three different reference time points.**

| **Beginning of S phase [s]** (134) | **End of S phase [s]** (134) | **NEBD [s]** | **Metaphase onset [s]** | **Anaphase onset [s]** | **Cytokinesis onset [s]** | **Chromosome decondensation [s]** |
|---|---|---|---|---|---|---|
| -900 | -330 | 0 | 145 | 180 | 280 | 300 |
| -1200 | -630 | -300 | -155 | -120 | -20 | 0 |
| -1180 | -610 | -280 | -135 | -100 | 0 | 20 |
